# Evolution of immune genes in island birds: reduction in population sizes can explain island syndrome

**DOI:** 10.1101/2021.11.21.469450

**Authors:** Mathilde Barthe, Claire Doutrelant, Rita Covas, Martim Melo, Juan Carlos Illera, Marie-Ka Tilak, Constance Colombier, Thibault Leroy, Claire Loiseau, Benoit Nabholz

**Author notes:** Last co-authors (Claire Loiseau, Benoit Nabholz).

## Abstract

Shared ecological conditions encountered by species that colonize islands often lead to the evolution of convergent phenotypes, commonly referred to as “island syndrome”. Reduced immune functions have been previously proposed to be part of this syndrome, as a consequence of the reduced diversity of pathogens on island ecosystems. According to this hypothesis, immune genes are expected to exhibit genomic signatures of relaxed selection pressure in island species. In this study, we used comparative genomic methods to study immune genes in island species (N = 20) and their mainland relatives (N = 14). We gathered public data as well as generated new data on innate (TLR: Toll-Like Receptors, BD: Beta Defensins) and acquired immune genes (MHC: Major Histocompatibility Complex classes I and II), but also on hundreds of genes with various immune functions. As a control, we used a set of 97 genes, not known to be involved in immune functions based on the literature, to account for the increased drift effects of the lower effective population sizes in island species. We used synonymous and non-synonymous variants to estimate the selection pressure acting on immune genes. We found that BDs and TLRs have higher ratios of non-synonymous over synonymous polymorphisms (Pn/Ps) than randomly selected control genes, suggesting that they evolve under a different selection regime. However, simulations show that this is unlikely to be explained by ongoing positive selection or balancing selection. For the MHC genes, which evolve under balancing selection, we used simulations to estimate the impact of population size variation. We found a significant effect of drift on immune genes of island species leading to a reduction in genetic diversity and efficacy of selection. However, the intensity of relaxed selection was not significantly different from control genes, except for MHC class II genes. These genes exhibit a significantly higher level of non-synonymous loss of polymorphism than expected assuming only drift and evolution under frequency dependent selection, possibly due to a reduction of extracellular parasite communities on islands. Overall, our results showed that demographic effects lead to a decrease in the immune functions of island species, but the relaxed selection that is expected to be caused by a reduced parasite pressure may only occur in some categories of immune genes.

## Introduction

Island colonizers face new communities of competitors, predators and parasites in a small area with limited resources, which generally results in high extinction rates of colonizers (Losos and Ricklefs, 2009). Oceanic island faunas are characterized by a low species richness, coupled with high population densities for each species (MacArthur and Wilson, 1967; Warren et al., 2015) - which translates into communities with, on average, lower levels of inter-specific interactions and higher levels of intra-specific competition (but see Rando et al., 2010 for an example of character displacement due to competition among island finch species). These shared island characteristics are thought to underlie the evolution of convergent phenotypes, in what is called the ‘island syndrome’ (Baeckens and Van Damme, 2020). Convergence has been documented in multiple traits, such as size modification (dwarfism or gigantism; Lomolino, 2005), reduction of dispersal (Baeckens and Van Damme, 2020), shift towards K life-history strategies (Boyce,1984; Covas, 2012; MacArthur and Wilson, 1967), evolution of generalist traits (Blondel, 2000; Warren et al., 2015), or changes in color and acoustic signals (Doutrelant et al., 2016; Grant, 1965).

Reduced immune function has also been hypothesized to be an island syndrome trait, directly linked to reduced parasite pressure on islands (Lobato et al., 2017; Matson and Beadell, 2010; Wikelski et al.,2004). Island parasite communities are i) less diverse (Beadell et al., 2006; Illera et al., 2015; Loiseau et al.,2017; Maria et al., 2009; Pérez-Rodríguez et al., 2013), and ii) could be less virulent due to the expansion of the ecological niche expected by the theory of island biogeography. In fact, island parasites are often more generalist than their mainland counterparts, which could lead to a reduced virulence due to the trade-off between replication capacity and resistance against host immune defenses (Garamszegi, 2006; Hochberg and Møller, 2001; Pérez-Rodríguez et al., 2013). Overall, a reduction of parasitic pressure should lead to a weakening of the immune system due to the costs of maintaining efficient immune functions (Lindström et al., 2004; Matson and Beadell, 2010; Wikelski et al., 2004). Such reduction may have important implications for the ability of these populations to resist or tolerate novel pathogens. The introduction of avian malaria in the Hawaiian archipelago, and the subsequent extinctions and population declines of many endemic species is the most emblematic example (Van Riper III et al., 1986; Wikelski et al., 2004).

Immunological parameters, such as blood leukocyte concentration, antibodies or other immune proteins (e.g. haptoglobin), hemolysis, and hemagglutination (Lee et al., 2006; Matson and Beadell, 2010) may serve as proxies to determine population immune functions. To date, the majority of studies that focused on island avifauna have found ambiguous results, with either no support for a reduced immune response on island species (Beadell et al., 2007; Matson, 2006), or contrasting results, such as a lower humoral component (total immunoglobulins) on islands, but a similar innate component (haptoglobin levels) between island and mainland species (Lobato et al., 2017). The use of immune parameters as proxies of immune function is fraught with difficulties (Lobato et al., 2017). The study of molecular evolution of immune genes therefore represents an alternative strategy to tackle this question. However, it is necessary to distinguish neutral effects (i.e. the demographic effects resulting from island colonization) from selective ones, the potential relaxation of selection pressures due to the changes in the pathogen community.

The bottleneck experienced by species during island colonization leads to a decrease in genetic variability (Frankham, 1997). A reduced genetic diversity at loci involved in immunity should have a direct implication on immune functions (Hale and Briskie, 2007 but see; Hawley et al., 2005; Spurgin et al.,2011). Also, small population sizes increase genetic drift, which may counteract the effect of natural selection on weakly deleterious mutations (Ohta, 1992). Several recent studies found a greater load of deleterious mutations in island species (Kutschera et al., 2020; Leroy et al., 2021b; Loire et al., 2013; Robinson et al., 2016; Rogers and Slatkin, 2017). Finally, it is necessary to differentiate genes involved in the innate versus the acquired immune response. The innate immune response is the first line of defense and is composed of phagocytes, macrophages and dendritic cells. These cells allow non-specific recognition of pathogens (Akira, 2003; Alberts et al., 2002). For example, Toll-Like Receptors (TLR; transmembrane proteins) trigger a chain reaction leading to the production of various substances, including antimicrobial peptides such as beta-defensins (BD) that have active properties in pathogen cell lysis (Velová et al., 2018). On the other hand, the acquired immune system allows a specific response, characterized by immune memory. Major Histocompatibility Complex (MHC) genes code for surface glycoproteins that bind to antigenic peptides, and present them to the cells of the immune system; class I and II genes ensure the presentation of a broad spectrum of intra- and extracellular-derived peptides, respectively (Klein, 1986). Although all these genes are directly involved in the identification and neutralization of pathogens, previous studies found that they evolve under different selection regimes: TLRs and BDs are under purifying selection which usually results in the selective removal of deleterious alleles and stabilizing selection (Grueber et al., 2014; van Dijk et al., 2008), whereas MHC genes are under balancing selection (Bernatchez and Landry, 2003).

Recent studies on birds (Gonzalez-Quevedo et al., 2015a, 2015b), amphibians (Belasen et al., 2019), and lizards (Santonastaso et al., 2017) found that the demographic history of island populations led to the loss of genetic variants at immune genes involved in pathogen recognition, such as TLRs and MHC. For example, Santonastaso et al., (2017) revealed that the polymorphism pattern in MHC genes and microsatellites covary positively with island area in *Podarcis* lizards, suggesting a dominant role for genetic drift in driving the evolution of the MHC. Gonzalez-Quevedo, et al. (2015a) found a similar pattern comparing TLR and microsatellite polymorphism in the Berthelot pipit, *Anthus berthelotii,* an endemic species from Macaronesia, supporting a predominant role of genetic drift in TLR evolution. However, these studies did not explicitly test the hypothesis of a relaxed selection pressure on islands imposed by an impoverished parasite community. All other things being equal, it is expected that the polymorphism of a coding sequence decreases with population size (Buffalo, 2021; Leroy et al., 2021b). Therefore, a decrease in polymorphism with population size could not be taken as a proof of a relaxation in the selection pressure.

To be able to demonstrate a change in natural selection, a traditional approach is to contrast polymorphism of synonymous sites (Ps) with polymorphism of non-synonymous sites (Pn). Synonymous mutations do not change amino acid sequences, whereas non-synonymous mutations do. Thus, synonymous mutations are expected to be neutral while non-synonymous could be subject to selection.

Following population genetic theory, in a diploid population, Ps = 4 *Ne μ* and Pn = 4 *Ne μ f*, where *Ne* is the effective population size, *μ* is the mutation rate and *f* is a function that integrates the probability of an allele to segregate at a given frequency. *f* depends on the distribution of the fitness effect (DFE) of mutations (Eyre-Walker and Keightley, 2007). This distribution scales with *Ne* as the fitness effect is dependent on *Ne* multiplied by the coefficient of selection *s* (Kimura, 1962). The nearly-neutral theory predicts that the DFE includes a large proportion of mutations with a *Ne*s* close to 0 (Ohta, 1992). As a consequence, an increase of *Ne* will lead to an increase of the fitness effect of weakly deleterious mutations, in such a way that these mutations will be more easily removed from the population by natural selection, therefore reducing Pn relative to Ps, leading to a negative correlation between Pn/Ps and Ps (through Ne; Welch et al., 2008). The presence of linked mutations, that are positively selected, does not change this relationship qualitatively (Castellano et al., 2018; Chen et al., 2020 and our simulations below).

Shifts in the parasitic community on islands are expected to have an impact on the Pn/Ps ratio of immune genes. However, the fixation probability depends on the product *Ne*s*, and variation in *Ne* is also expected to impact the efficacy of selection and thus the Pn/Ps ratio across the entire transcriptome, particularly in the presence of slightly deleterious mutations (Charlesworth and Eyre-Walker, 2008; Leroy et al., 2021b; Loire et al., 2013; Ohta, 1992). In addition, due to their lower population sizes, island birds compared to continental species exhibit a genome-wide reduction in genetic diversity and efficacy of selection (Kutschera et al., 2020; Leroy et al., 2021b). Therefore, we expect a similar reduction in immune genes’ diversity even without any change in the parasite pressure.

To disentangle the effect of population size from a change in parasite pressure and estimate the impact of demography on the efficacy of selection, we studied a dataset of 34 bird species (20 insular and 14 mainland species; Figure 1) combining the 24 species of Leroy et al. (2021b) and 10 newly generated by targeted-capture sequencing (Table 1). We randomly selected protein-coding genes (i.e., control genes) involved in various biological functions (Fijarczyk et al., 2016; Leroy et al., 2021b). The selection pressure acting on the randomly selected control genes is expected to be similar between island and mainland bird species. Therefore, the variation of Pn/Ps of the control genes is only dependent on the variation of *Ne*. In contrast, if a reduced parasite pressure on islands directly impacts the evolution of immune genes, the Pn/Ps of immune genes is expected to show a larger variation between island and continental species than the control genes. More specifically, for genes under purifying selection, non-synonymous weakly deleterious mutations, normally eliminated under strong selection, would be maintained, leading to an increase of Pn/Ps. By contrast, for genes under balancing selection, non-synonymous advantageous mutations, normally maintained in the polymorphism under strong selection, would be fixed or eliminated leading to a decrease of Pn/Ps (Figure 2).

**Figure 1:**
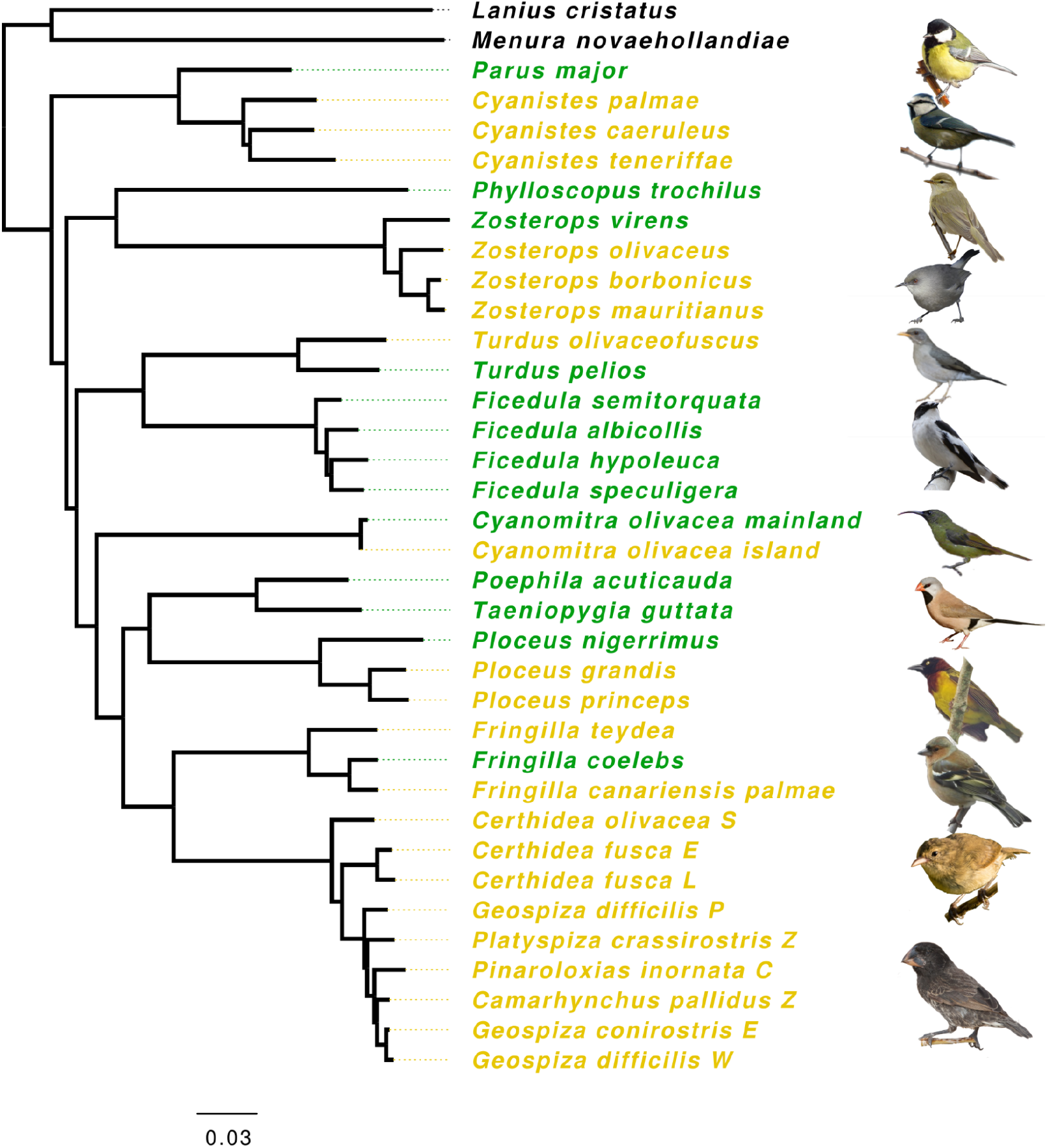
Phylogeny based on mitochondrial genes of species from the dataset reconstructed by maximum likelihood method (IQTREE model GTR+Gamma). Species names in yellow indicate island species, and in green, mainland species. Ultrafast bootstrap values are provided in the supplementary methods. Some relationships are poorly supported. Bird representations are not to scale. Photos from top to bottom: *P. major, C. caeruleus, P. trochilus, Z. borbonicus, T. pelios, F. albicollis, C. olivacea, P. acuticauda, P. grandis, F. coelebs, C. fusca, G. conirostris*. Photo credits: A. Chudý, F. Desmoulins, E. Giacone, G. Lasley, Lianaj, Y. Lyubchenko, B. Nabholz, J.D. Reynolds, K. Samodurov, A. Sarkisyan, Wimvz, Birdpics, T. Aronson, G. Lasley, P. Vos (iNaturalist.org); M. Gabrielli (*Zosterops borbonicus*).

**Figure 2:**
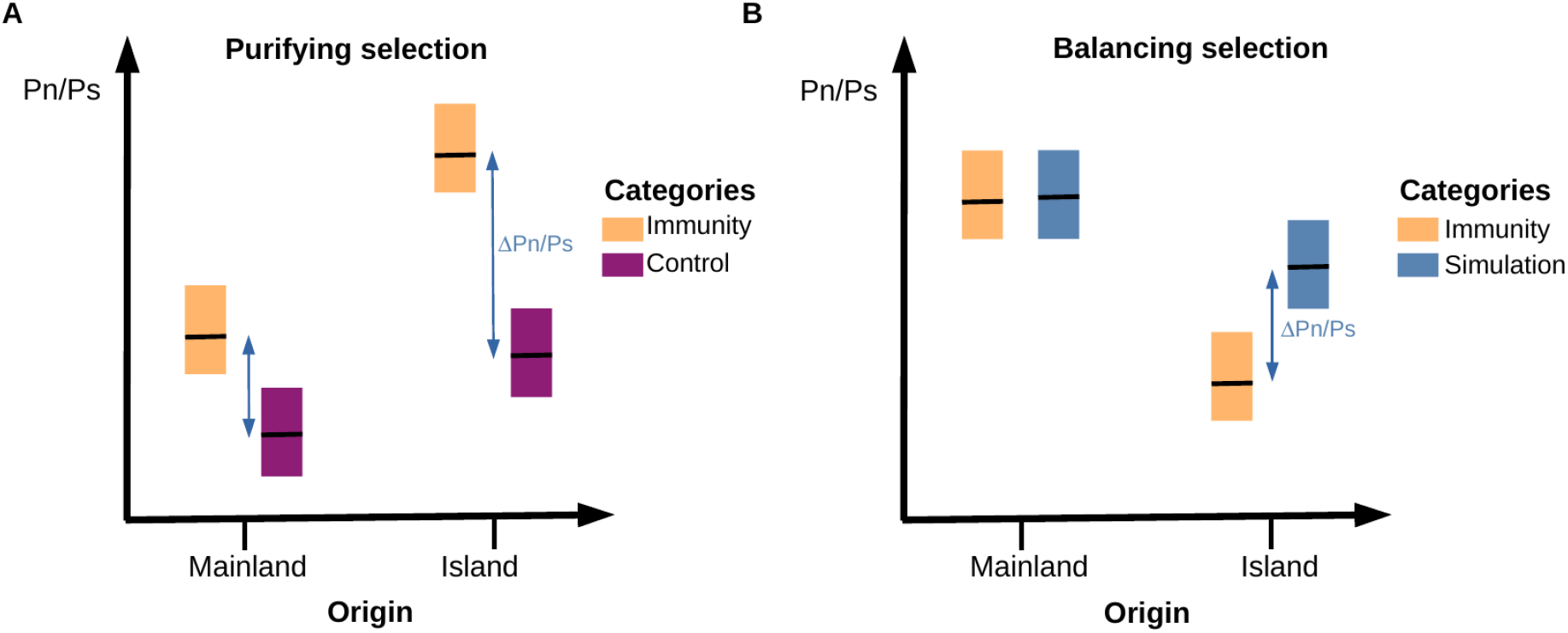
Conceptual diagram showing the expected results under the hypothesis of a relaxation in the selection pressure of the immune genes in island species due to a change in the parasitic community. A) Genes evolving under purifying selection where control genes are randomly selected protein-coding genes. B) Genes evolving under balancing selection where controls are obtained from SLiM simulations of genes evolving under the same balancing selection but different population size. Under the hypothesis of a relaxed selection as a consequence of the reduced diversity of pathogens on island ecosystems, the difference in Pn/Ps between categories (ΔPn/Ps) is expected to be different between species’ origin, leading to a statistical interaction between gene categories and origin.

**Table 1:**
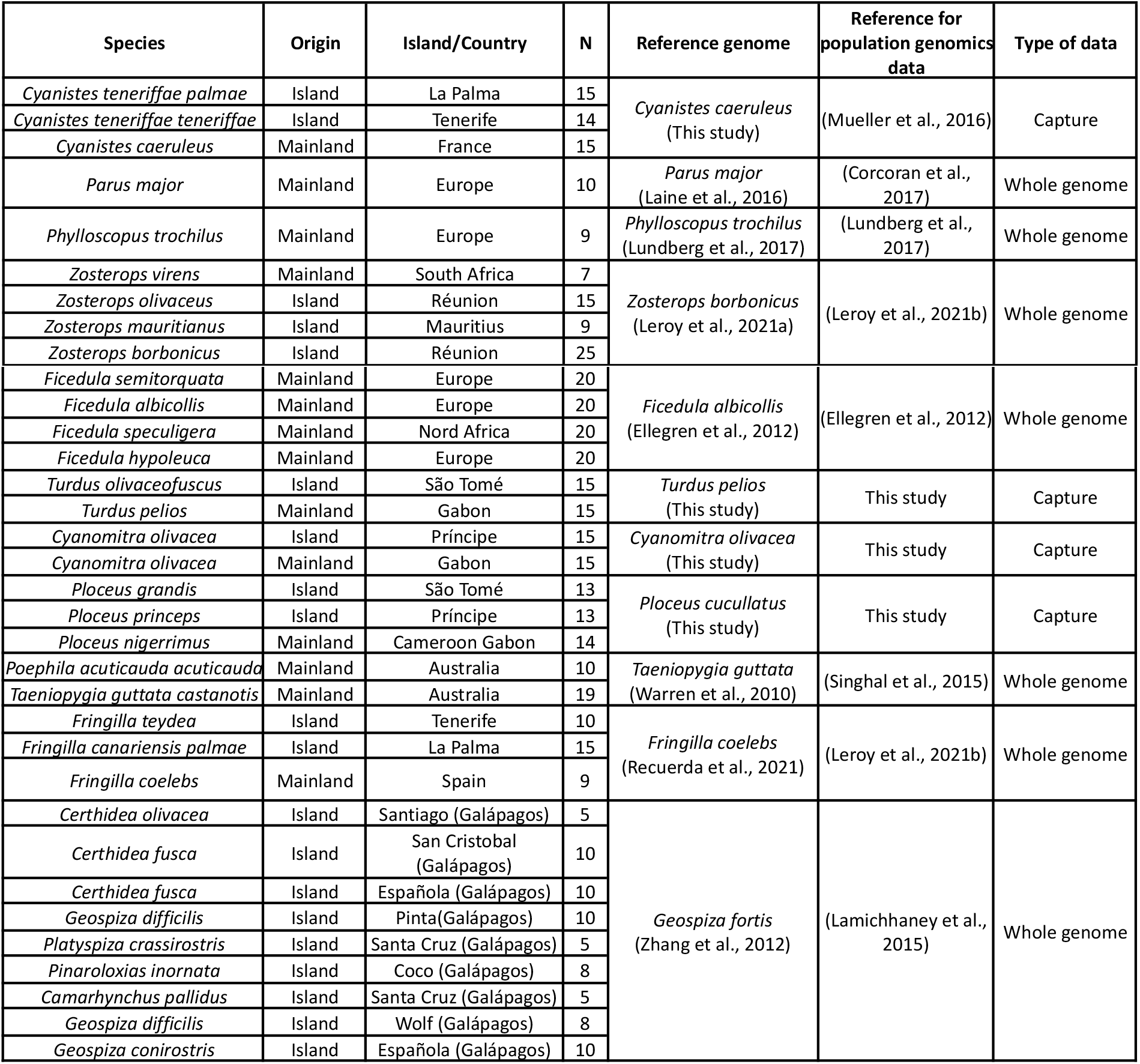
List of species and sampling localities, along with the type of data obtained and the number of individuals (N).

## Methods

### Dataset

Alignments of Coding DNA Sequences (CDS) of individuals from 24 species were obtained from Leroy et al. (2021b). In addition, data for ten other species (six and four from islands and mainland, respectively) were newly generated for this study by targeted-capture sequencing. Blood samples and subsequent DNA extractions were performed by different research teams. The complete dataset consisted of 34 bird species (20 and 14 insular and mainland species respectively; Table 1; Figure 1). We filtered alignments in order to retain only files containing a minimum of five diploid individuals per site (Table 1).

Sequence enrichment was performed using MYBaits Custom Target Capture Kit targeting 21 immune genes: 10 Toll-Like receptors (TLR), 9 Beta Defensins (BD), 2 Major Histocompatibility Complex (MHC) and 97 control genes (see below). We followed the manufacturer’s protocol (Rohland and Reich, 2012). Illumina high-throughput sequencing, using a paired-end 150 bp strategy, was performed by Novogene (Cambridge, UK).

#### Newly generated draft genome sequence

We generated whole genome sequences at moderate coverage (~40X) for *Turdus pelios, Ploceus cucullatus* and *Cyanomitra olivacea* (from Gabon). Library preparation from blood DNA samples and Illumina high-throughput sequencing using a paired-end 150 bp strategy were performed at Novogene (Cambridge, UK). Raw reads were cleaned using FastP (vers. 0.20.0; Chen et al., 2018). Genomes assemblies were performed using SOAPdenovo (vers. 2.04) and Gapcloser (v1.10) (Luo et al., 2012) with parameters “-d 1 -D 2” and a kmers size of 33. Protein annotation was performed by homology detection using genBlastG (She et al., 2011; http://genome.sfu.ca/genblast/download.html) and the transcriptome of the collared flycatcher (*Ficedula albicollis*; assembly FicAlb1.5; Ellegren et al., 2012) as reference.

#### Capture data processing

Reads from targeted-capture sequencing were cleaned with FastP (vers. 0.20.0; Chen et al., 2018). Reads of each individual were mapped respectively to the nearest available reference genomes using bwa mem (vers. 0.7.17; Li, 2013; Table 1), with default parameters. Samtools (vers. 1.3.1; Li et al., 2009) and Picard (vers. 1.4.2; Picard Toolkit 2019) were used to convert the mapping files, order and index reads according to their position on the chromosomes (or scaffolds) of the reference genomes or on the draft genomes generated in this study for *Ploceus, Cyanomitra* and *Turdus.* Duplicate reads were marked using MarkDuplicates (vers. 1.140; Picard Toolkit 2019). SNP calling was performed with Freebayes (vers. 1.3.1; Garrison and Marth, 2012). Freebayes output file (VCF file) was converted to a fasta file by filtering out sites with a minimum quality of 40 and a sequencing depth between 10 and 1000X (sites outside these thresholds were treated as missing data, i.e., ‘N’). CDS were then extracted from the alignments using the coordinates of the annotations (gff files). CDS were aligned using MACSE (vers. 2.03; Ranwez et al., 2011) to prevent frameshift mutation errors and GNU-parallel (Tange, 2018) was used to parallelise the computation.

#### Selection and identification of immune and control genes

We defined several groups of immune genes to compare with the control genes. The control group consisted of 97 protein-coding genes randomly selected in the genome of *Zosterops borbonicus* (Leroy et al., 2021a). These control genes allowed the estimation of the average selection pressure that a gene, not involved in the immune response, undergoes in the genome under a given effective population size. These genes were single copy (absence of paralogue) and had a variable GC content representative of the whole transcriptome.

For the immune genes, we selected three sets of genes from i) a limited set of genes (Core Group) where functions are unambiguously related to immunity, and ii) two larger sets of genes (Database-group & Sma3s-group), obtained through an automatic annotation pipeline.

The Core Group included MHC class I and class II genes, 10 Toll-Like Receptors (TLRs; Velová et al.,2018) and 9 Beta Defensins (BD; Chapman et al., 2016). The Database group included genes identified by Immunome Knowledge Base (Ortutay and Vihinen, 2009, http://structure.bmc.lu.se/idbase/IKB/; last access 04/02/2020) and InnateDB (Breuer et al., 2013, http://www.innatedb.com; last access 04/02/2020). We also added a set of genes for which the genetic ontology indicated a role in immune functions. To do so, we used the chicken *(Gallus gallus)* annotation (assembly GRCg6a downloaded from Ensembl database in March 2020; https://www.ensembl.org/). We identified genes with the terms “immun*” or “pathogen*” in their Gene Ontology identifiers description (directory obtained from http://geneontology.org/). This set included 2605 genes considered to be involved in immunity, although some may be only indirectly involved in immunity or have a small impact on immune functions. Finally, the third set of genes (Sma3s-group) has been built up through the Sma3s-group program (vers. 2; Munoz-Mérida et al., 2014). This program annotated sequences in order to be associated with biological functions through gene ontology identifiers. The annotation of the genome of *F. albicollis* allowed us to identify 3136 genes associated with the genetic ontology “immune system processes”. Like for the Database group, this set may include genes with various functions in the immune response. It should be noted that Sma3s-group and Database-group were not mutually exclusive, and some genes were present in both groups. An analysis was performed to identify and exclude genes under balancing selection from Database-group and Sma3s-group sets using BetaScan (vers. 2; Siewert and Voight, 2020), due to the potentially antagonistic responses of these genes. Very few genes (only 2 and 3 genes from Database-group and Sma3s-group sets) were identified and removed from the analysis (see Detection of genes under balancing selection in Supplementary Methods).

#### Test for contamination and population structure

We used the program CroCo (vers. 1.1; Simion et al., 2018) to identify candidates for cross-species contamination (see supplementary materials for details). Overall, we did not detect a clear case of cross-species contamination in our dataset (Figure S1; Barthe and Nabholz, 2022). Contigs identified as potential contamination always involve a pair of species belonging to the same genus. In this case, contamination could be difficult to identify due to the low genetic divergence between species.

For the newly sequenced species, we also performed PCA analyses using allele frequencies of control genes. We used the function dudi.pca of adegenet R package (Jombart and Ahmed, 2011). This analysis aims to check for population structure and to detect potentially problematic individuals (i.e., contaminated individuals). This analysis led to the exclusion of 4 individuals *(Ploceus princeps* P6-174; *P. grandis* ST10_094; *P. nigerrimus* G3_016; *C. teneriffae* TF57) for which we suspected contamination. Otherwise, no extra population structure was detected (Figure S2-S4; Barthe and Nabholz, 2022).

#### Hidden paralogy

We computed the statistic F_IS_ = 1-H_0_/H_e_ where H_0_ is the average number of heterozygous individuals observed (H_0_ = #heterozygous / n; where n is the sample size) and H_e_ is the expected number of heterozygous individuals at Hardy-Weinberg (HW) equilibrium (H_e_ = (n/(n-1) 2 * p * (1-p))*n where n is the sample size and p the allele frequency of a randomly chosen allele). F_IS_ varies between −1 and 1 with positive value representing excess of homozygous individuals and negative value representing excess of heterozygous individuals compared to the HW proportions. Gene with high value of nucleotide diversity (Pi) and negative value of F_IS_ could represent a potential case where hidden paralogous sequences have not been separated and where all the individuals present heterozygous sites in the positions where a substitution occurred between the paralogous copies. Five sequences corresponding to the TLR21 genes appeared problematic (Pi > 0.01 and F_IS_ < −0.5; Figure S5; Barthe and Nabholz, 2022) and were excluded from further analyses.

The MHC genes were more difficult to analyse. Indeed, heterozygosity could be comparable to divergence under balancing selection. This made the identification of orthologs very difficult. We identified a variable number of genes among species (from 1 to 10 genes for MHC class I and MHC class II). We checked the sequence similarity for the 10 copies of the MHC class II in *F. albicollis* and the 7 copies of the MHC class I genes in *C. caeruleus* using cd-hit (Fu et al., 2012). For MHC class II, sequence divergences were always higher than 15% indicating that reads were likely correctly assigned to their corresponding gene copy. For MHC class I, sequence similarity could be as high as 95%. In this case, we relied on the fact that the reads from very similar paralogous copies were not be confidently assigned to a gene copy sequence by the mapping software. This should lead to a low mapping score quality and were likely to be discarded during the genotype calling procedure. For example, 3 out of 7 of the *Cyanistes* MHC class I genes were not correctly genotyped and were missing from our final dataset.

### Data Analysis

#### SLiM simulations

We used SLiM (vers. 3.3.2; Haller and Messer, 2017) to estimate the impact of demographic changes on polymorphism patterns under various selection regimes. The following parameters were used in all simulations. Sequences of 30kb with a mutation rate of 4.6e^−9^ substitutions/site/generation were simulated (Smeds et al., 2016). Recombination was set to be equal to mutation rate. Introns/exons pattern was reproduced by simulating fragments of 3kb separated by one bp with a very high recombination rate of 0.1 rec./site/generation. We chose 3kb because TLR CDS were typically single-exon sequences of 2-3kb (Velová et al., 2018). Five types of mutations were possible: i) neutral synonymous mutations, ii) codominant non-synonymous mutations with a Distribution of Fitness Effect (DFE) following a gamma law of mean = −0.025 and shape = 0.3, which corresponds to the DFE estimated in Passerines by Rousselle et al. (2020), iii) codominant non-synonymous mutations positively selected with s = 0.1, iv) non-synonymous mutations under balancing selection with an effect on fitness initially set at 0.01 but re-estimated by the program at each generation according to the mutation frequency in the population, thus including a frequency-dependent effect and v) non-synonymous mutations under overdominance with a dominance coefficient of 1.2.

We simulated a coding sequence organization where positions one and two of the codons were considered as non-degenerated sites, with the non-synonymous types of mutations previously described were possible in various proportions. The third position was considered as completely neutral where only synonymous mutations could appear.

In the absence of control genes evolving under balancing selection, we used SLiM to generate a set of control genes for this category. We simulated two populations of 270,000 and 110,000 individuals, representing mainland and island effective population size respectively.

We also explored the effect of positive and balancing selection on the pattern of Ps and Pn/Ps in a population of size 50,000, 110,000, 270,000 and 500,000. In order to speed up the computational time, we reduced the population size by a factor 100 and rescaled mutation rate, recombination rate and selection coefficient accordingly running 10 replicates per simulation.

All the details of the simulation parameters, calculations of non-synonymous polymorphism rate (Pn) and synonymous polymorphism rate (Ps) of simulated sequences, as well as SLiM command lines are provided in Supplementary Methods and Materials.

#### Polymorphism analyses

Synonymous (Ps) and non-synonymous (Pn) nucleotide diversities were estimated from seq_stat_coding written from the Bio++ library (Available as Supplementary data; Guéguen et al., 2013). The mean Pn/Ps was computed as the sum of Pn over the sum of Ps (Wolf et al., 2009). Ps of concatenated sequences of control genes were estimated for each species of our dataset. For the whole-genome sequenced species, we compared the Pn/Ps and Ps estimated from the 97 control genes with the values from Leroy et al., (2021b; ~5000 genes used in their study). Pn/Ps and Ps correlations showed a R^2^ of 0.6 and 0.95 respectively (Figure S6; Barthe and Nabholz, 2022). Thus, the 97 control genes used in our study were representative of the larger set of genes from Leroy et al (2021b). This allowed us to identify *Phylloscopus trochilus* as an outlier. Unlike for all other species (e.g. *Fringilla coelebs,* Figure S7; Barthe and Nabholz, 2022), synonymous polymorphism level was correlated to the amount of missing data in *P. trochilus* alignments (Figure S7; Barthe and Nabholz, 2022). As such, we excluded *P. trochilus* from further analysis.

The mean Pn/Ps, calculated from the concatenated sequences of genes from the same gene class (control genes; BD; TLR; MHC I; MHC II; Database-group; Sma3s-group), was estimated for each bird species. Alternative transcripts were identified based on the genomic position in the GFF file. If several transcripts were available, one transcript was randomly selected. Pn/Ps estimates based on less than four polymorphic sites were excluded from the analysis, as were those with no polymorphic non-synonymous sites.

#### Statistical analyses

To estimate the impact of demographic history on genome-wide polymorphism of island species and the potentially reduced constraints on their immune genes, we computed the ratio of non-synonymous nucleotide diversity over synonymous nucleotide diversity (Pn/Ps). A linear mixed model was performed, using the Pn/Ps ratio as dependent variable and, as explanatory variables, the mainland or insular origin of species as well as the category of genes (packages lme4 and lmerTest (Bates et al., 2012; Kuznetsova et al., 2017)). In order to take into account the phylogenetic effect, the taxonomic rank “family”was included as a random effect in the model. We also used a generalized linear mixed model (using the function glmer of the package lme4) with the family “Gamma(link=“log”)” which led to the same results (Figure S15 to S24; Barthe and Nabholz, 2022). Five linear mixed models were defined i) model including origin and gene category parameters and also the interaction effect ii) model using both origin and gene category parameters, iii) model with only the gene category parameter, iv) model with only the origin parameter, and finally v) null model. In some cases, the phylogenetic effect was difficult to estimate because the number of species per family was reduced to one. In that case, we choose to reduce the number of families by grouping Turdidae with Muscicapidae, Nectariniidae, and Estrildidae with Ploceidae and Fringillidae within Thraupidae. The results obtained with these family groupings were similar to the original model (Table S1; Barthe and Nabholz, 2022), except when stated. The categories Database-group and Sma3s-group were tested separately from the Core group because they contained hundreds of genes annotated using the automatic pipeline that were only available for species with genome wide data. Database-group and Sma3s-group were not analysed simultaneously because they contained a partially overlapping set of genes. Finally, genes evolving under purifying selection and genes evolving under balancing selection were also analysed separately. Model selection was based on two methods. First, we used the difference in corrected Akaike Information Criterion (ΔAICc) calculated using the qpcR package (Spiess and Spiess, 2018). Second, a model simplification using an ANOVA between models was also performed.

We also tested an alternative model using the difference between Pn/Ps of immune genes and control genes (ΔPn/Ps) as dependent variable, and species origin as explanatory variable. Under the hypothesis of a relaxation in selection pressure on islands due to a change in the parasite community, we expected the ΔPn/Ps to be higher on island species compared to the mainland ones and, therefore, the species origin (i.e., mainland or island) to be significant. In this model, we used the Phylogenetic Generalized Least Squares model (PGLS; implemented in the “nlme” packages; Pinheiro et al., 2017). This model assumed that the covariance between species follows a Brownian motion evolution process along the phylogeny (implemented using the “corBrownian” function from the ‘“ape”’ package; Paradis and Schliep, 2019). The species phylogeny was estimated using mitochondrial genes and a maximum likelihood inference implemented in IQTREE (model GTR+Gamma and ultrafast bootstrap; Nguyen et al., 2014; median of 11,134 bp analysed per species). The phylogeny with the bootstrap support is provided as supplementary material.

All the statistical analyses were performed using R (R Core Team, 2018), and dplyr package (Wickham,2016). Graphical representations were done using ggplot2, ggrepel, ggpubr and ggpmisc (Aphalo, 2020; Kassambara, 2018; Slowikowski et al., 2018; Wickham, 2016).

## Results

For the 150 individuals (10 species with 15 individuals each) for which we generated new data by targeted capture sequencing, an average of 3.3 million paired-ends reads per individual was generated (Table S1; Barthe and Nabholz, 2022). Additionally, we generated three new draft assemblies using 40x pair-end illumina data for the species for which no closely related reference genomes were available. N50 and total size were 1.11 Gb and 27.9 kb for *Cyanomitra olivaceus*; 1.10 Gb and 31.7 kb for *Ploceus cucullatus* and 1.13 Gb and 14.3 kb for *Turdus pelios*. After mapping, genotyping and cleaning, we analysed 86 control and 16 immune genes on average per species, out of the 141 targeted genes (120 control and 21 immune related genes; Table S4; Barthe and Nabholz, 2022). For the species with whole-genome sequences, we analysed 106 control and 20 immune genes on average per species, out of the 141 targeted genes, and 875 and 688 genes on average in the Database-group and Sma3s-group respectively (Table S4; Barthe and Nabholz, 2022).

For the species for which full genome sequences were available, the Ps and Pn/Ps estimated using the control genes reflect the Ps and Pn/Ps of the whole transcriptome (Figure S6; Barthe and Nabholz, 2022).

### Population genetics of BD and TLR immune genes

In order to characterize the selection regimes shaping the BD and TLR polymorphisms (Figure 3), we first analyzed the variation of Pn/Ps ratios among gene categories using a linear mixed model.

**Figure 3:**
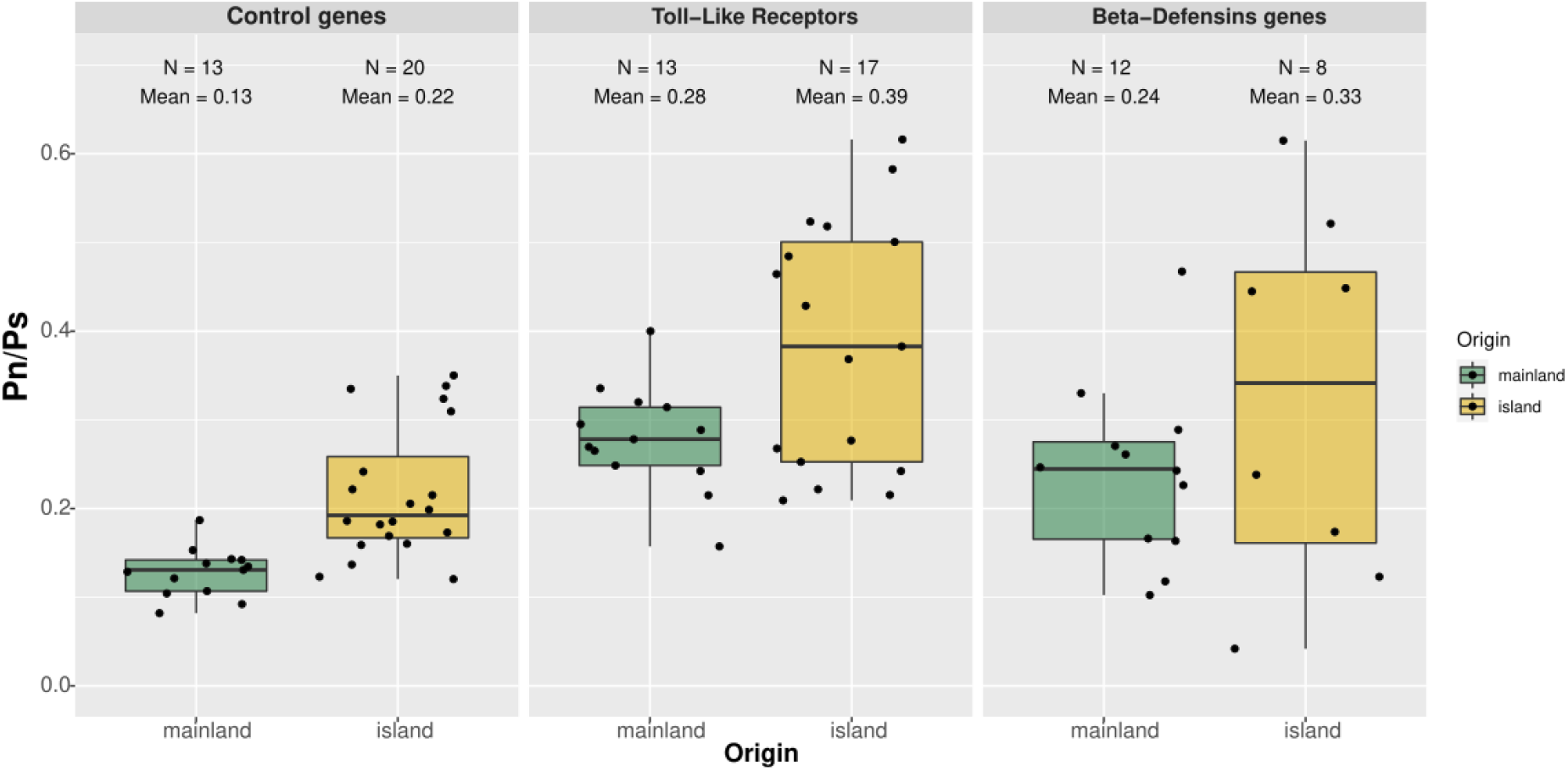
Pn/Ps according to species origin (mainland in green and insular in orange) for different gene categories under purifying selection. The number of species (N), and the mean Pn/Ps are shown for each modality.

Model selection based on AICc as well as model selection approach based on simplification with ANOVA identified the model n° 2, including the origin (i.e., mainland or island) and gene category without interaction (Table 2). In this model, island origin of species is associated with a greater Pn/Ps (0.14 vs. 0.10; Table 3; p < 0.01). Gene categories corresponding to TLRs and BDs showed a significantly higher Pn/Ps than control genes (Table 3; p < 0.001). Our statistical analysis confirmed that island birds have a higher Pn/Ps ratio than mainland relatives, in agreement with the nearly-neutral theory of evolution. It also reveals that immune genes have a higher Pn/Ps than randomly selected control genes suggesting that BD and TLR evolve under a different selection regime than non-immune related genes.

**Table 2:** Statistical model explaining Pn/Ps variation of Toll-Like Receptors, Beta-Defensins genes, and control genes. The p-values of ANOVA test between simpler models are not reported if a more complex model explains a larger proportion of the variance.

| n° | Model | Model selection by AIC |  |  | ANOVA test |  |  |  |
| --- | --- | --- | --- | --- | --- | --- | --- | --- |
| | | AICc | ΔAICc | Likelihood | n° 1 | 2 | 3 | 4 |
| 1 | Pn/Ps~ 1+ category +origin+ category *origin | -5.39 | 8.83 | 0.01 |  | 0.63 |  |  |
| 2 | Pn/Ps~ 1+ category +origin | -14.22 | 0 | 1 |  |  | 0.002 | 3.71E-05 |
| 3 | Pn/Ps~ 1+ category | -11.8 | 2.42 | 0.3 |  |  |  |  |
| 4 | Pn/Ps~ 1+ origin | -6.83 | 7.39 | 0.02 |  |  |  |  |
| 5 | Pn/Ps~ 1 | -6.44 | 7.78 | 0.02 |  |  |  |  |

**Table 3:**
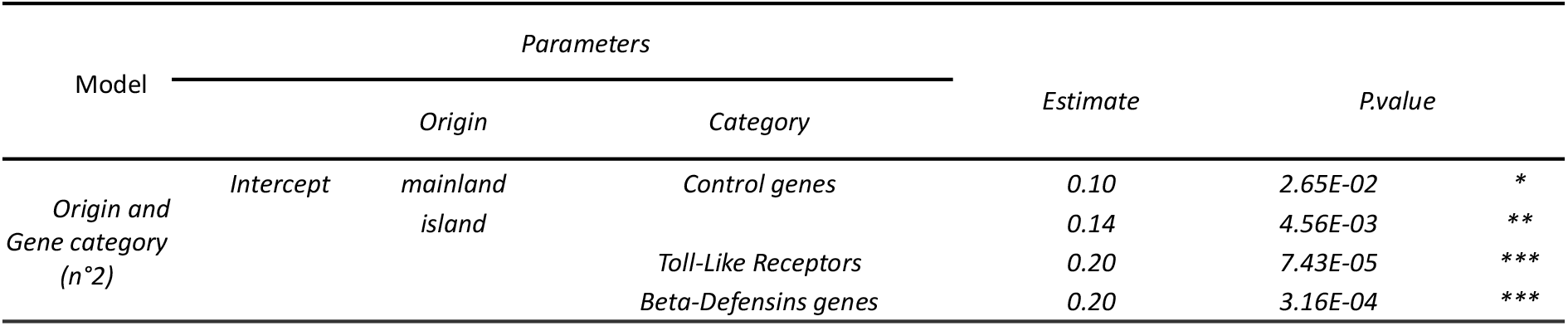
Summary of the model n°2, best statistical model selected using AICc explaining variation in Pn/Ps in control genes, Toll-Like receptors and Beta-Defensins genes under purifying selection with origin, gene category parameters. * indicates significant values: * < 0.05; ** < 0.01; *** < 0.001.

Next, we investigated the cause of the higher Pn/Ps of immune genes by testing three hypotheses. First, we excluded a bias due to a lower number of immune genes, and therefore higher variance in the estimation of Pn/Ps in immune genes. Immune genes still had significantly higher Pn/Ps compared to a random subsample of control genes of comparable size (Figure S8 & S9; Barthe and Nabholz, 2022). Second, the Pn/Ps of immune genes could be inflated by positive selection. It is well known that immune genes are subject to frequent adaptation due to arms race evolution with pathogens (Enard et al., 2016; Shultz and Sackton, 2019; Velová et al., 2018). We evaluated the effect of positively selected genes on the Pn/Ps using SLiM simulations with both positively and negatively selected mutations. The presence of recurrent positive selection could increase the Pn/Ps leading to a higher Pn/Ps in immune genes if this category was more prone to adaptive evolution (Figure 4A). However, positive selection always led to a drastic decrease in Ps due to genetic sweep effect at linked sites (Figure 4B). BDs and TLRs had a slightly higher or similar Ps than control genes (Figure S9, mean Ps = 0.007, 0.004 and 0.003 for BDs, TLRs and control genes respectively, effect of gene category p < 0.1; Barthe and Nabholz, 2022) and, as a consequence, even if positive selection is likely to have impacted the evolution of immune genes, it is not the cause of the higher Pn/Ps observed here. Third, balancing selection could be present, at least temporarily, in the evolution of BDs and TLRs genes (Kloch et al., 2018; Levy et al., 2020). Simulation analyses confirmed that balancing selection causes an increase of Ps and Pn/Ps (Figure 4C & 4D). However, a change in effective population size had an opposite effect on the Pn/Ps according to whether selection was negative or balancing. In the presence of slightly deleterious mutations, Pn/Ps decreases with *Ne* whereas it increases in the presence of balancing selection. Island birds had higher Pn/Ps ratios than mainland birds for BDs and TLRs. Therefore, we can rule out balancing selection as the main factor explaining the high Pn/Ps of immune genes because, in this case, Pn/Ps of island birds should be lower. Another possible explanation is a relaxed selection of immune genes. It is likely that immune genes are overall less constrained than the control genes. It has been shown that evolutionary constraints are more related to gene expression than to function (Drummond et al., 2005; Drummond and Wilke, 2008) and therefore, functionally important genes could still have a high Pn/Ps.

**Figure 4:**
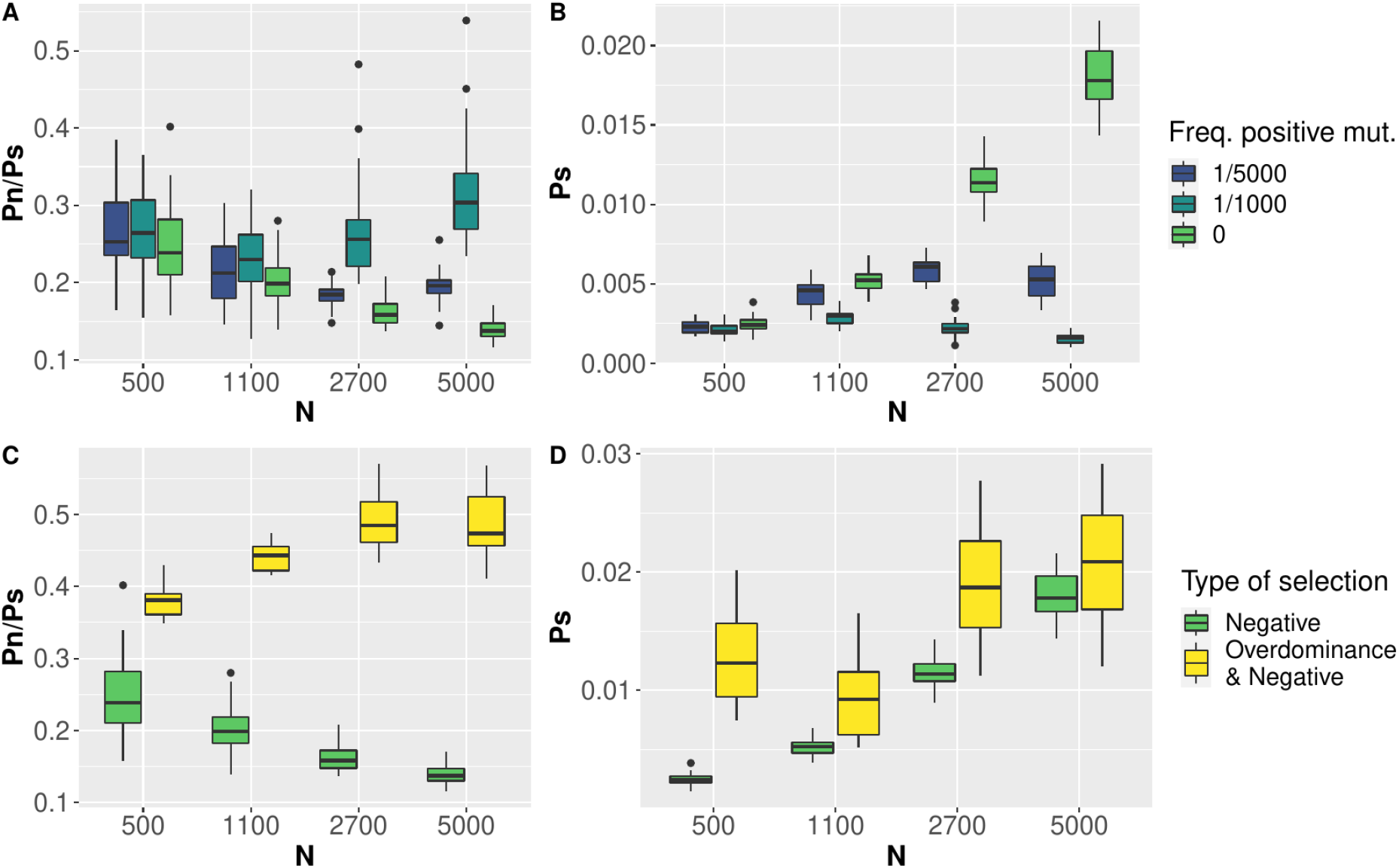
Neutral polymorphism (Ps) and ratio of selected over neutral polymorphism (Pn/Ps) estimated from SLiM simulations. A) Pn/Ps as a function of population size, N and B) Ps as a function of N. In both A and B, color indicates the frequency of positively selected mutations compared to deleterious mutations. C) Pn/Ps as a function of N and D) Ps as a function of N. In both C and D, yellow indicates simulations with overdominance mutation (h = 1.2) and negatively selected mutations and green indicates simulations with only negatively selected mutations.

Overall, our analyses do not support a strong impact of ongoing adaptive mutation or balancing selection on BDs and TLRs. However, these immune genes do not evolve as random genes (not involved in immune functions) and present a significantly higher Pn/Ps of 0.20 (p < 0.001; Table 3).

### No evidence of a reduced impact of the parasite communities on the polymorphism pattern of immunes genes in island birds

For BDs and TLRs, the best model selected includes the origin (i.e., mainland or island) and gene category without interaction, corresponding to model n°2 (see above and Table 2). This model has no interaction between origin and gene categories invalidating the hypothesis of a reduced parasite communities on islands (Figure 2).

For larger sets of genes, identified using an automatic pipeline and gene annotation, model selection based on AICc and simplification with ANOVA (Table S5, S8; Barthe and Nabholz, 2022) identified models n°4 that included origine parameters which associated a higher Pn/Ps of at least 0.07 for island species (p < 0.001; Table S6, S7, S9, S10; Barthe and Nabholz, 2022, Figure 5). Model selection by simplification with ANOVA identified models n°1 with interaction effect between origin and gene category associated with a reduced Pn/Ps for TLR and BD genes of island species that invalidate our hypothesis (Table S7, S10; Barthe and Nabholz, 2022).

The alternative statistical approach using the difference between Pn/Ps of immune genes and control genes (ΔPn/Ps) as dependent variable, and species origin as explanatory variable under a PGLS framework lead to similar results. Island was never associated to a statistically higher ΔPn/Ps (Table S2; Barthe and Nabholz, 2022) providing no support for an increased relaxed selection of immune genes in island species.

**Figure 5:**
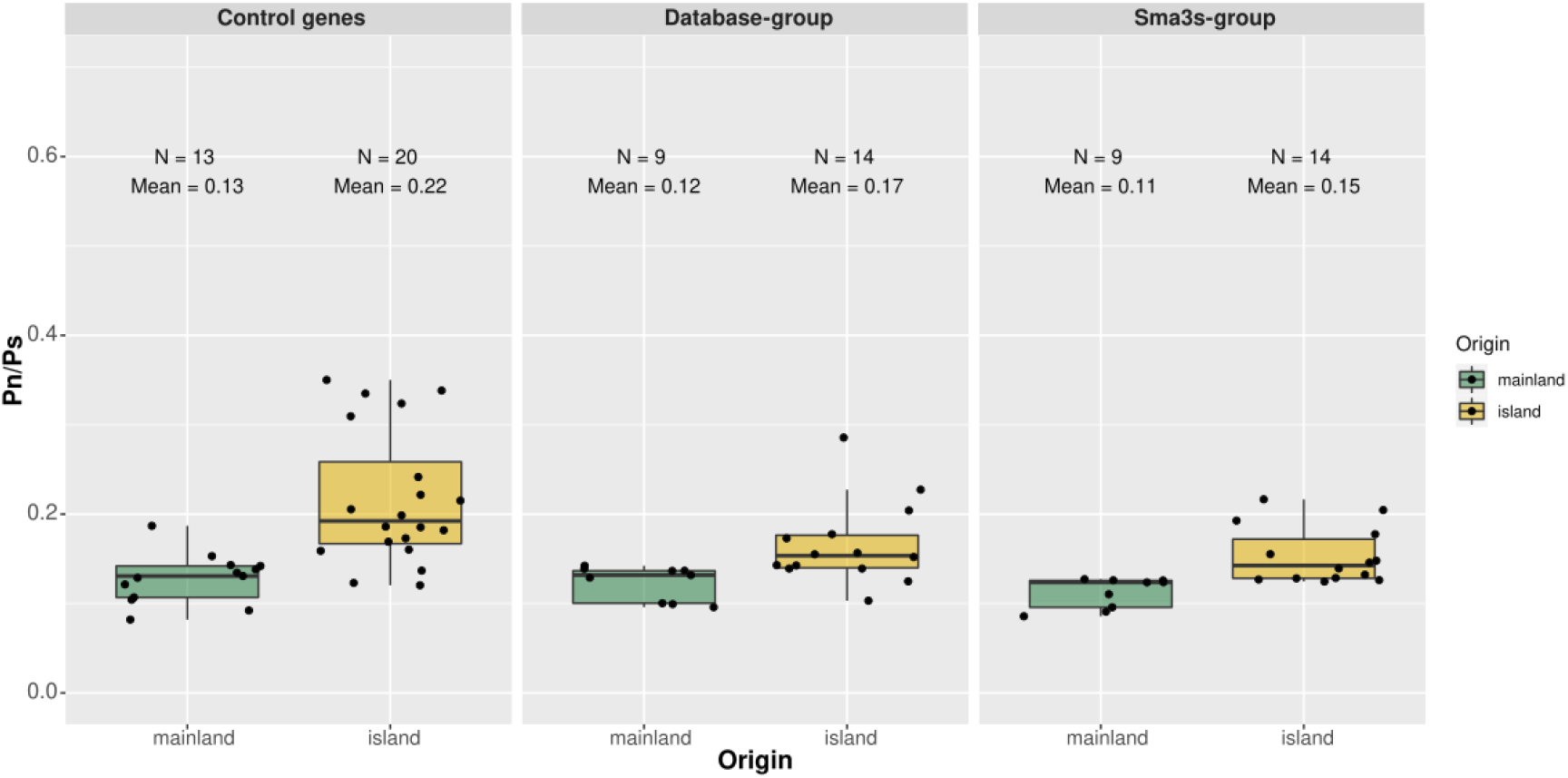
Boxplot of Pn/Ps according to species origin (mainland in green and insular in orange) for different gene categories under purifying selection. The number of individuals (N), and the mean Pn/Ps are shown for each modality.

### Genes under balancing selection

First, we estimated the effect of population size variation on the Pn/Ps of the genes evolving under balancing selection by simulating sequences under frequency dependent or overdominance selection using SLiM (see Methods and Supplementary Methods). The simulation under frequency dependent selection revealed an average Pn/Ps equal to 0.8 for island species and 1.2 for mainland species (Figure 6). Under overdominance, simulated sequences from island and mainland populations respectively have an average Pn/Ps equal to 0.54 and 1.03 (Figure 6).

**Figure 6:**
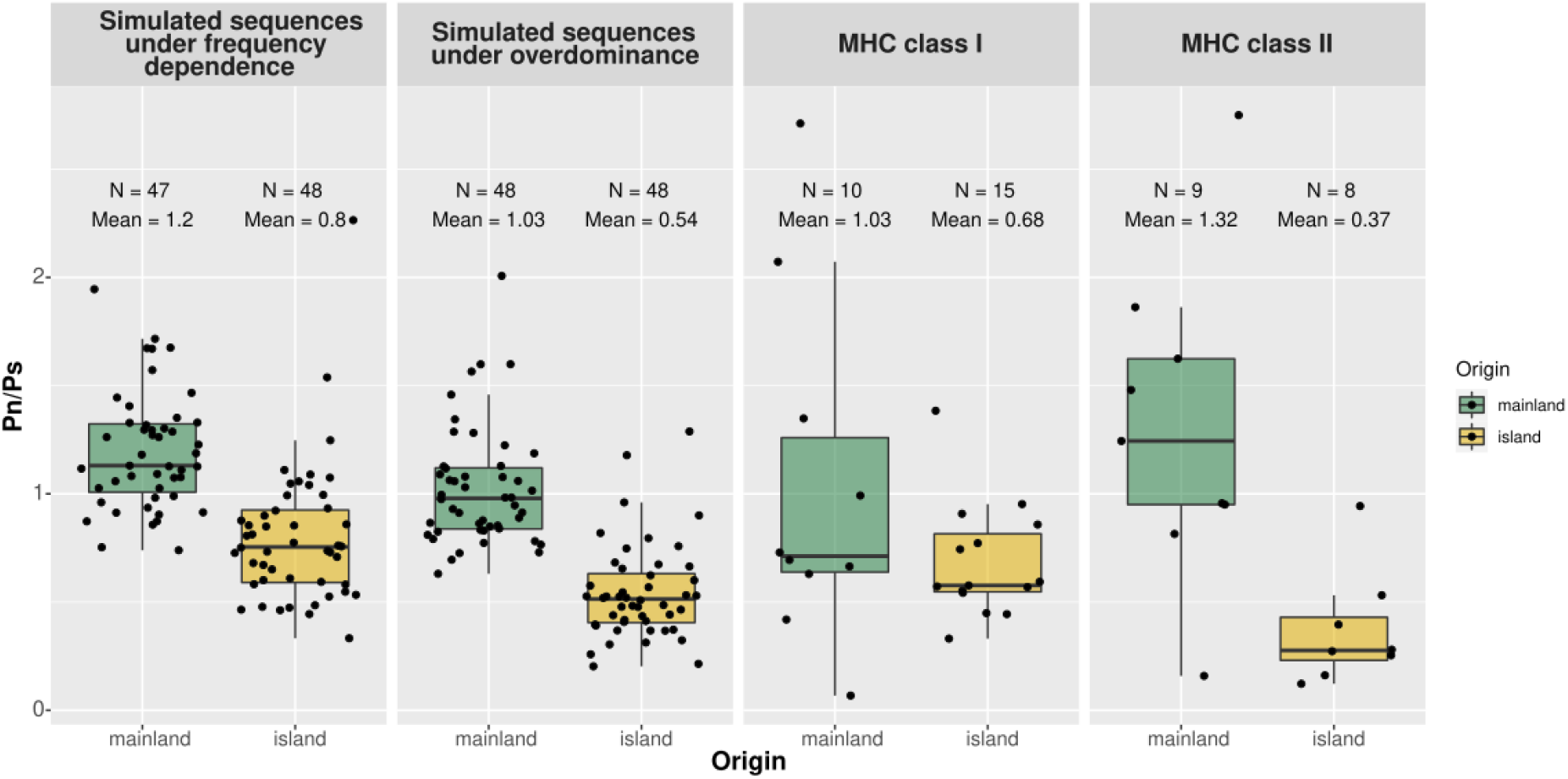
Boxplot of Pn/Ps according to species origin (mainland in green and insular in orange) for different gene categories under balancing selection. The number of species (N), and the mean Pn/Ps are shown for each modality. The control groups correspond to the results obtained from simulated sequence via SLiM (see Methods and Supplementary Methods Simulation of control genes under balancing selection).

Using simulations under frequency dependent selection as well as simulations under the overdominance, model selection by AIC identifies the model n°4 with origin, contrary to the method by simplification with ANOVA which identified the full model (model n°1) therefore including significant interaction between origin and genes category (Table 4). This interaction effect is significant for the MHC II (p < 0.05, Table S12; Barthe and Nabholz, 2022) but not for MHC I. As expected, island species have a significantly lower Pn/Ps in MHC genes compared to mainland species (p < 0.01; except for the full model based on control genes evolving under overdominance Table S12; Barthe and Nabholz, 2022).

**Table 4:**
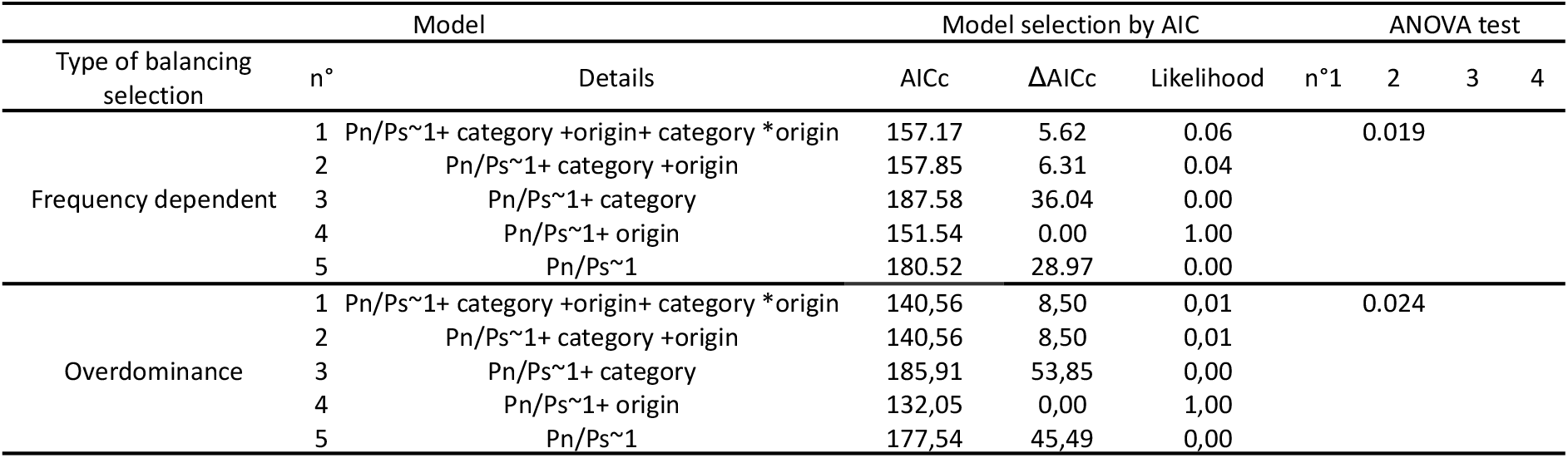
Statistical model explaining Pn/Ps variation of genes under balancing selection (i.e MHC class I and II), and simulated sequences under i) frequency dependent or ii) overdominance. The p-values of ANOVA test between simpler models are not reported if a more complex model explains a larger proportion of the variance.

## Discussion

On oceanic islands, the depauperate parasite community is expected to lead to a relaxation of selection on the immune system. In this study, we found support for such an effect, but only on MHC class II genes and using simulated sequences under balancing selection as control. No effect was detected for MHC class I genes nor for innate immune genes (TLRs and BDs), evolving under purifying selection. On these sets of genes, increased drift effects on island populations limit the efficacy of selection in accordance with the nearly-neutral theory (Ohta, 1992). The ability to distinguish between the selective and nearly-neutral processes (relaxed selection due to environmental change vs. drift) could only be achieved by our approach of using random genes (i.e., “control genes”) to estimate the genome-wide effect of potential variation in effective population size between populations.

### Effects of effective population size variation

Our results support the nearly-neutral theory of evolution for those genes under purifying selection, whereby strong genetic drift acting on small island populations reduces the efficacy of natural selection, leading to an increase in non-synonymous nucleotide diversity compared to the mostly neutral, synonymous nucleotide diversity (i.e., Pn/Ps; Ohta, 1992). This is materialized by a genome-wide increase in frequency of weakly deleterious mutations (Kutschera et al., 2020; Leroy et al., 2021b; Loire et al.,2013; Robinson et al., 2016; Rogers and Slatkin, 2017).

For genes evolving under balancing selection, we performed simulations under the hypotheses of overdominance (heterozygote advantage) or frequency-dependent (rare allele advantage). Our results showed reduced Pn/Ps for smaller population sizes (Figure 6, S10, S11). This simulation confirmed our expectations (Figure 2) that a reduction in the efficacy of selection results in a decrease in the frequency of non-synonymous polymorphism, as, under normal circumstances, selection maintains those mutations at intermediate frequencies. It also matches what we obtained for the empirical results, where both MHC classes I and II had a reduced Pn/Ps in island birds. This result supports that the fitness effect of having non-synonymous polymorphisms segregating at high frequencies is not strong enough to counteract entirely the effect of genetic drift on islands.

### Effects of selection on immune genes

For immune genes, we tried to characterize the nature of the selection acting on BDs and TLRs genes. Comparing those genes with control genes and using simulations, we were able to rule out that directional positive selection and balancing selection had a major impact shaping the polymorphism of these immune genes. In contrast, the pattern of Pn/Ps between island and mainland populations is in line with the effect of purifying selection in the presence of slightly deleterious mutations. However, no effect was detected on insular species, beyond what could be attributed to genetic drift. This is in line with the result of Gonzalez-Quevedo et al. (2015b) and Grueber et al. (2013) who found that TLR genetic diversity was mostly influenced by genetic drift. At first sight, this result seems not in line with the fact that island parasite communities are less diverse (Beadell et al., 2006; Loiseau et al., 2017; Maria et al., 2009; Pérez-Rodríguez et al., 2013; but see Illera et al., 2015). However, a reduced number of pathogens has also been found to be associated with a higher prevalence in birds and reptiles from the Macaronesian archipelago (Illera and Perera, 2020). Therefore, these two patterns, i.e. a less diverse pathogen’s community on islands with a higher prevalence, could still imply a strong selection pressure on immune genes.

In contrast, for MHC genes that unambiguously evolve under balancing selection, MHC class II genes presented a reduction in non-synonymous polymorphism larger than the effects of drift alone, when simulated sequences are used as control. This was the only case where a role for relaxed selection pressures in the molecular evolution of immune genes could be invoked.

Our results are in accordance with the hypothesis of Lee (2006), which proposes that innate and acquired immunity may exhibit distinct responses to changes in pressures due to different costs and benefits. However, they contrast with the study of Santonastaso et al. (2017) that identified no change in selection pressures on MHC II genes in a lizard species and concluded that their evolution was mostly governed by drift. Similarly, Agudo et al. (2011) also found a prominent role for genetic drift over selection in the evolution of MHC II genes in the Egyption vulture (*Neophron percnopterus*).

Our results rely on simulations that may be affected by the choice of the parameter values. First, we performed simulations using a fixed effective population size (*Ne*) estimated from the polymorphism data. Using others values of *Ne* had a weak impact on the relative difference between island and mainland species for the overdominance type of selection (Figure S10, S11; Barthe and Nabholz, 2022). Secondly, we simulated two types of selection, namely overdominance (Doherty and Zinkernagel, 1975) and frequency-dependent (Slade and McCallum, 1992), but it has been argued that the maintenance of MHC polymorphism could be the result of fluctuating selection (Hill, 1991). Additionally, recombination has also been put forward as a mechanism responsible for generating diversity (Spurgin et al., 2011). Therefore, our results for the MHC II genes, which is based on the relative difference between Pn/Ps of island and mainland species comparing empirical and simulated data, should be taken cautiously as their significance can be dependent on the specific parameters that we used, although we did our best to select a realistic range of parameters.

The observed difference between MHC class I and II could be explained by their different pathogen targets: MHC class I genes are primarily involved in the recognition of intracellular pathogens (Kappes and Strominger, 1988), while MHC class II genes are directly involved in the recognition of extracellular pathogens (Bjorkman and Parham, 1990). These differences could lead to variable selection pressures depending on the extracellular versus intracellular parasite communities present on islands. In addition, the relaxed selection pressures on MHC II genes from adaptive immunity is in line with a reduction in acquired immunity parameters found by Lobato et al. (2017).

Future work should take into account that there is an extensive variation in the number of MHC gene copies across the avian phylogeny (Minias et al., 2019; O’Connor et al., 2020). Particularly, it was recently discovered that Passerines have a very dynamic evolution of duplication/loss events compared to other birds (Minias et al., 2019). Here, we used the two copies of MHC gene I and II currently annotated in the collared flycatcher genome as target sequences for our targeted-capture sequencing. The future improvement of genome assembly, resulting from the development of long-reads technology (Peona et al., 2021, 2018), should help to annotate with increased precision all MHC copies and to study the whole repertoire of MHC genes.

### Consequences of drift and selection on immunity

The potential relaxation of the natural selection acting on immune genes in island species is expected to reduce immune functions and increase susceptibility of island populations to pathogens. This is true even if this relaxation is only the consequence of a reduction in the effective population size and not caused by a reduction of the pressure exerted by the parasitic community. This is in line with the results of Hawley et al. (2005) and Belasen et al. (2019) who showed that a decrease in diversity of immune loci (MHC II or through immune proxy) was associated with a reduction in immune functions. It should be noted that even if migration rate is reduced on islands, sedentary and endemic island species are not completely free from the exposure of exogen pathogens through migratory birds (Levin et al., 2013).

As a final remark, we would like to stress that more research is needed (i) to ascertain both selection pressures on innate and adaptive immune responses and the load of deleterious mutations due to drift, also identified by an increasing body of work (Loire et al., 2013; Robinson et al., 2016; Rogers and Slatkin,2017; Kutschera et al., 2020; Leroy et al., 2021b), and (ii) to better describe island parasite communities. To date, most of the studies investigated intracellular parasite communities on islands, and more specifically haemosporidian parasites, avian pox and coccidian parasites (Cornuault et al., 2012; Illera et al., 2015, 2008; Ishtiaq et al., 2010; Loiseau et al., 2017; Martinez et al., 2015; Padilla et al., 2017; Pérez-Rodríguez et al., 2013;Silva-Iturriza et al., 2012), whereas very few evaluated the extracellular parasite diversity, such as helminths (Nieberding et al., 2006, but see the review of Illera and Perera 2020 for reptiles). Metabarcoding of parasites is a new technique to evaluate at the same time both communities of intracellular and extracellular parasites (Bourret et al., 2021) and might therefore be a promising approach to compare their communities in island and mainland populations.

## Conclusion

Our comparative population genomics study has investigated the combined effects of drift and selection on immune genes from island and mainland passerines. The study of synonymous and non-synonymous polymorphism of these genes confirmed that island species, with smaller population sizes than their mainland counterparts, were more impacted by drift, which induces a load of weakly deleterious mutations in their genome. Indeed most of the genes studied here involved in the immune response do not show a statistically different pattern from control genes. Only MHC II genes, involved in the recognition of extracellular pathogens, showed a reduction in their non-synonymous polymorphism in island species. This response, which may be attributed to reduced selection pressures on these genes, could be associated with the suspected reduced parasitic communities on islands. The increased load of deleterious mutations as well as the potential relaxed selection pressures on MHC II support the reduced immune functions of island species, which could be added to the list of other convergent responses of the island syndrome.

## Supporting information

https://doi.org/10.6084/m9.figshare.16954921.v7

## Acknowledgements

In Gabon, we thank the Director and the guides of the Lekedi Park, Marie Charpentier for her help in organizing the expedition, and Elisa Lobato and Alexandre Vaz for fieldwork assistance and outreach work. In São Tomé and Príncipe, we thank the Directorate of the Environment and the Department for Nature Conservation, its directors—Arlindo Carvalho and Victor Bonfim—Guilhermino, the Association Monte Pico, its president Luis Mário, and its members. Elisa Lobato, Philippe Perret, Octávio Veiga, Bikegila, and Yelli provided invaluable assistance in the field. Permissions for fieldwork were given by the authorities of São Tomé and Príncipe and Gabon (CENAREST authorization No. AR0053/12/MENESTFPRSCJS/ CENAREST/CG/CST/CSAR). Permits for the Canary Islands were provided by the Regional Government (Ref.: 2012/0710), and the Cabildo of La Palma and Tenerife. In Montpellier, we thank the blue tit team (https://oreme.org/observation/ecopop/mesanges/) for the capture of the individuals used in this study. The analyses benefited from the Montpellier Bioinformatics Biodiversity (MBB) platform services. This research was conducted in the scope of the international twin-lab “LIA – Biodiversity and Evolution” between CIBIO (Portugal) and ISEM and CEFE-CNRS (France). This is ISEM publication n° ISEM 2022-223. Preprint version 4 of this article has been peer-reviewed and recommended by Peer Community In Evolutionary Biology (https://doi.org/10.24072/pci.evolbiol.100153)

## Data, scripts, code, and supplementary information availability

Datasets, scripts, supplementary figures and texts are available on figshare (Barthe and Nabholz,2022). The reads newly generated for this study have been deposited in the NCBI Sequence Read Archive under the bioproject PRJNA724656.

## References

Agudo R, Alcaide M, Rico C, Lemus JA, Blanco G, Hiraldo F, Donázar JA (2011) Major histocompatibility complex variation in insular populations of the Egyptian vulture: inferences about the roles of genetic drift and selection. Molecular Ecology 20, 2329–2340. https://doi.org/10.1111/j.1365-294X.2011.05107.x.

Akira S (2003) Toll-like receptor signaling. Journal of Biological Chemistry 278, 38105–38108. https://doi.org/10.1074/jbc.R300028200.

Alberts B, Johnson A, Lewis J, Raff M, Roberts K, Walter P (2002) Innate immunity. Molecular Biology of the Cell. https://doi.org/10.1093%2Faob%2Fmcg023

Aphalo PJ (2020) ggpmisc: Miscellaneous Extensions to “ggplot2”(R package version 0.3. 6).

Baeckens S, Van Damme R (2020) The island syndrome. Current Biology 30, R338–R339. https://doi.org/10.1016/j.cub.2020.03.029

Barthe M, Doutrelant C, Covas R, Melo M, Illera JC, Tilak M-K, Colombier C, Leroy T, Loiseau C, Nabholz B (2022) Evolution of immune genes in island birds: reduction in population sizes can explain island syndrome. bioRxiv, 2021.11.21.469450, ver. 4 peer-reviewed and recommended by Peer Community in Evolutionary Biology. https://doi.org/10.1101/2021.11.21.469450

Barthe M, Nabholz B (2022) Supplementary materials for “Evolution of immune genes in island birds: reduction in population sizes can explain island syndrome”. Figshare. https://doi.org/10.6084/m9.figshare.16954921.v7

Bates DM, Maechler M, Bolker B, Walker S (2012) Package ‘lme4’. CRAN R Found Stat Comput.

Beadell JS, Atkins C, Cashion E, Jonker M, Fleischer RC (2007) Immunological change in a parasite-impoverished environment: divergent signals from four island taxa. PLoS One 2:e896,. https://doi.org/10.1371/journal.pone.0000896.

Beadell JS, Ishtiaq F, Covas R, Melo M, Warren BH, Atkinson CT, Bensch S, Graves GR, Jhala YV, Peirce MA (2006) Global phylogeographic limits of Hawaii’s avian malaria. Proceedings of the Royal Society B: Biological Sciences 273, 2935–2944. https://doi.org/10.1098/rspb.2006.3671.

Belasen AM, Bletz MC, S LD, Toledo LF, James TY (2019) Long-term habitat fragmentation is associated with reduced MHC IIB diversity and increased infections in amphibian hosts. Frontiers in Ecology and Evolution 6,. https://doi.org/10.3389/fevo.2018.00236.

Bernatchez L, Landry C (2003) MHC studies in nonmodel vertebrates: what have we learned about natural selection in 15 years? Journal of Evolutionary Biology 16, 363–377. https://doi.org/10.1046/j.1420-9101.2003.00531.x.

Bjorkman PJ, Parham P (1990) Structure, function, and diversity of class I major histocompatibility complex molecules. Annual Review of Biochemistry 59, 253–288. https://doi.org/10.1146/annurev.bi.59.070190.001345.

Blondel J (2000) Evolution and ecology of birds on islands: trends and prospects. Vie et Milieu/Life & Environment 205–220. https://hal.sorbonne-universite.fr/hal-03186916

Bourret V, Gutiérrez López R, Melo M, Loiseau C (2021) Metabarcoding options to study eukaryotic endoparasites of birds. Ecology and Evolution 11, 10821–10833. https://doi.org/10.1002/ece3.7748.

Boyce MS (1984) Restitution of gamma-and k-selection as a model of density-dependent natural selection. Annual Review of Ecology and Systematics 15, 427–447. https://doi.org/10.1146/annurev.es.15.110184.002235

Breuer K, Foroushani AK, Laird MR, Chen C, Sribnaia A, Lo R, Winsor GL, Hancock RE, Brinkman FS, Lynn DJ (2013) InnateDB: systems biology of innate immunity and beyond—recent updates and continuing curation. Nucleic Acids Research 41:D1228–D1233,. https://doi.org/10.1093/nar/gks1147.

Buffalo V (2021) Quantifying the relationship between genetic diversity and population size suggests natural selection cannot explain Lewontin’s paradox (G Sella, Ed.). ELife 10, e67509. https://doi.org/10.7554/eLife.67509.

Castellano D, James J, Eyre-Walker A (2018) Nearly neutral evolution across the Drosophila melanogaster genome. Molecular Biology and Evolution 35, 2685–2694. https://doi.org/10.1093/molbev/msy164

Chapman H JR, O H, AS K, RH C, RL W, J. (2016) The evolution of innate immune genes: purifying and balancing selection on β-defensins in waterfowl. Molecular Biology and Evolution 33, 3075–3087. https://doi.org/10.1093/molbev/msw167.

Charlesworth J, Eyre-Walker A (2008) The McDonald–Kreitman test and slightly deleterious mutations. Molecular Biology and Evolution 25, 1007–1015. https://doi.org/10.1093/molbev/msn005.

Chen J, Glémin S, Lascoux M (2020) From drift to draft: how much do beneficial mutations actually contribute to predictions of Ohta’s slightly deleterious model of molecular evolution? Genetics 214, 1005–1018. https://doi.org/10.1534/genetics.119.302869.

Chen S, Zhou Y, Chen Y, Gu J (2018) fastp: an ultra-fast all-in-one FASTQ preprocessor. Bioinformatics 34:i884–i890,. https://doi.org/10.1093/bioinformatics/bty560.

Corcoran P, Gossmann TI, Barton HJ, Slate J, Zeng K (2017) Determinants of the Efficacy of Natural Selection on Coding and Noncoding Variability in Two Passerine Species. Genome Biology and Evolution 9, 2987–3007. https://doi.org/10.1093/gbe/evx213.

Cornuault J, Bataillard A, Warren BH, Lootvoet A, Mirleau P, Duval T, Milá B, Thébaud C, Heeb P (2012) The role of immigration and in-situ radiation in explaining blood parasite assemblages in an island bird clade. Molecular Ecology 21, 1438–1452. https://doi.org/10.1111/j.1365-294X.2012.05483.x.

Covas R (2012) Evolution of reproductive life histories in island birds worldwide. Proceedings of the Royal Society B: Biological Sciences 279, 1531–1537. https://doi.org/10.1098/rspb.2011.1785.

van Dijk A, Veldhuizen EJ, Haagsman HP (2008) Avian defensins. Veterinary Immunology and Immunopathology 124, 1–18. https://doi.org/10.1016/j.vetimm.2007.12.006.

Doherty PC, Zinkernagel RM (1975) Enhanced immunological surveillance in mice heterozygous at the H-2 gene complex. Nature 256, 50–52. https://doi.org/10.1038/256050a0.

Doutrelant C, Paquet M, Renoult JP, Grégoire A, Crochet P-A, Covas R (2016) Worldwide patterns of bird colouration on islands. Ecology Letters 19, 537–545. https://doi.org/10.1111/ele.12588.

Drummond DA, Bloom JD, Adami C, Wilke CO, Arnold FH (2005) Why highly expressed proteins evolve slowly. Proceedings of the National Academy of Sciences 102, 14338–14343. https://doi.org/10.1073/pnas.0504070102.

Drummond DA, Wilke CO (2008) Mistranslation-induced protein misfolding as a dominant constraint on coding-sequence evolution. Cell 134, 341–352. https://doi.org/10.1016/j.cell.2008.05.042.

Ellegren H, Smeds L, Burri R, Olason PI, Backström N, Kawakami T, Künstner A, Mäkinen H, Nadachowska-Brzyska K, Qvarnström A (2012) The genomic landscape of species divergence in Ficedula flycatchers. Nature 491, 756–760. https://doi.org/10.1038/nature11584.

Enard D, Cai L, Gwennap C, Petrov DA (2016) Viruses are a dominant driver of protein adaptation in mammals. Elife 5, e12469. https://doi.org/10.7554/eLife.12469.

Eyre-Walker A, Keightley PD (2007) The distribution of fitness effects of new mutations. Nature Reviews Genetics 8, 610–618. https://doi.org/10.1038/nrg2146.

Fijarczyk A, Dudek K, Babik W (2016) Selective Landscapes in newt Immune Genes Inferred from Patterns of Nucleotide Variation. Genome Biology and Evolution 8, 3417–3432. https://doi.org/10.1093/gbe/evw236.

Frankham R (1997) Do island populations have less genetic variation than mainland populations? Heredity 78, 311–327. https://doi.org/10.1038/hdy.1997.46.

Fu L, Niu B, Zhu Z, Wu S, Li W (2012) CD-HIT: accelerated for clustering the next-generation sequencing data. Bioinformatics 28, 3150–3152. https://doi.org/10.1093/bioinformatics/bts565.

Garamszegi LZ (2006) The evolution of virulence and host specialization in malaria parasites of primates. Ecology Letters 9, 933–940. https://doi.org/10.1111/j.1461-0248.2006.00936.x.

Garrison E, Marth G (2012) Haplotype-based variant detection from short-read sequencing. ArXiv Preprint arXiv:1207.3907. https://doi.org/10.48550/arXiv.1207.3907.

Gonzalez-Quevedo C, Phillips KP, Spurgin LG, Richardson DS (2015) 454 screening of individual MHC variation in an endemic island passerine. Immunogenetics 67, 149–162. https://doi.org/10.1007/s00251-014-0822-1.

Gonzalez-Quevedo C, Spurgin LG, Illera JC, Richardson DS (2015) Drift, not selection, shapes toll-like receptor variation among oceanic island populations. Molecular Ecology 24, 5852–5863. https://doi.org/10.1111/mec.13437.

Grant PR (1965) The adaptive significance of some size trends in island birds. Evolution 355–367. https://doi.org/10.2307/2406446.

Grueber CE, Wallis GP, Jamieson IG (2013) Genetic drift outweighs natural selection at toll-like receptor (TLR) immunity loci in a re-introduced population of a threatened species. Molecular Ecology 22, 4470–4482. https://doi.org/10.1111/mec.12404.

Grueber CE, Wallis GP, Jamieson IG (2014) Episodic positive selection in the evolution of avian toll-like receptor innate immunity genes. PloS One 9, e89632. https://doi.org/10.1371/journal.pone.0089632.

Guéguen L, Gaillard S, Boussau B, Gouy M, Groussin M, Rochette NC, Bigot T, Fournier D, Pouyet F, Cahais V, Bernard A, Scornavacca C, Nabholz B, Haudry A, Dachary L, Galtier N, Belkhir K, Dutheil JY (2013) Bio++: Efficient Extensible Libraries and Tools for Computational Molecular Evolution. Molecular Biology and Evolution 30, 1745–1750. https://doi.org/10.1093/molbev/mst097.

Hale KA, Briskie JV (2007) Decreased immunocompetence in a severely bottlenecked population of an endemic New Zealand bird. Animal Conservation 10, 2–10. https://doi.org/10.1111/j.1469-1795.2006.00059.x.

Haller BC, Messer PW (2017) SLiM 2: Flexible, interactive forward genetic simulations. Molecular Biology and Evolution 34, 230–240. https://doi.org/10.1093/molbev/msw211.

Hawley DM, Sydenstricker KV, Kollias GV, Dhondt AA (2005) Genetic diversity predicts pathogen resistance and cell-mediated immunocompetence in house finches. Biology Letters 1, 326–329. https://doi.org/10.1098/rsbl.2005.0303.

Hill AV (1991) HLA associations with malaria in Africa: some implications for MHC evolution. ‘Mol. Evol. Major Histocompat. Complex’. pp. 403–420. (Springer: Berlin, Heidelberg) https://doi.org/10.1007/978-3-642-84622-9_33.

Hochberg ME, Møller AP (2001) Insularity and adaptation in coupled victim–enemy associations. Journal of Evolutionary Biology 14, 539–551. https://doi.org/10.1046/j.1420-9101.2001.00312.x.

Illera JC, Emerson BC, Richardson DS (2008) Genetic characterization, distribution and prevalence of avian pox and avian malaria in the Berthelot’s pipit (Anthus berthelotii) in Macaronesia. Parasitology Research 103, 1435–1443. https://doi.org/10.1007/s00436-008-1153-7.

Illera JC, Fernández-Álvarez Á, Hernández-Flores CN, Foronda P (2015) Unforeseen biogeographical patterns in a multiple parasite system in Macaronesia. Journal of Biogeography 42, 1858–1870. https://doi.org/10.1111/jbi.12548.

Illera JC, Perera A (2020) Where are we in the host-parasite relationships of native land vertebrates in Macaronesia? Ecosistemas 29, 1971. https://doi.org/10.7818/ECOS.1971

Institute B (2019) “Picard Toolkit”, Broad institute, GitHub repository. Picard Toolkit.

Ishtiaq F, Clegg SM, Phillimore AB, Black RA, Owens IP, Sheldon BC (2010) Biogeographical patterns of blood parasite lineage diversity in avian hosts from southern Melanesian islands. Journal of Biogeography 37, 120–132. https://doi.org/10.1111/j.1365-2699.2009.02189.x.

Jombart T, Ahmed I (2011) adegenet 1.3-1: new tools for the analysis of genome-wide SNP data. Bioinformatics 27, 3070–3071. https://doi.org/10.1093/bioinformatics/btr521.

Kappes D, Strominger JL (1988) Human class II major histocompatibility complex genes and proteins. Annual Review of Biochemistry 57, 991–1028. https://doi.org/10.1146/annurev.bi.57.070188.005015.

Kassambara A (2018) ggpubr:”ggplot2” based publication ready plots. R Package Version 01, 7.

Kimura M (1962) On the Probability of Fixation of Mutant Genes in a Population. Genetics 47, 713–719. https://doi.org/10.1093%2Fgenetics%2F47.6.713.

Klein J (1986) ‘Natural history of the major histocompatibility complex.’ (Wiley)

Kloch A, Wenzel MA, Laetsch DR, Michalski O, Bajer A, Behnke JM, Welc-Falęciak R, Piertney SB (2018) Signatures of balancing selection in toll-like receptor (TLRs) genes–novel insights from a free-living rodent. Scientific Reports 8, 1–10. https://doi.org/10.1038/s41598-018-26672-2.

Kutschera VE, Poelstra JW, Botero-Castro F, Dussex N, Gemmell N, Hunt GR, Ritchie MG, Rutz C, Wiberg RAW, Wolf JBW (2020) Purifying Selection in Corvids Is Less Efficient on Islands. Molecular Biology and Evolution. https://doi.org/10.1093/molbev/msz233.

Kuznetsova A, Brockhoff PB, Christensen RH (2017) lmerTest package: tests in linear mixed effects models. Journal of Statistical Software 82, 1–26. https://doi.org/10.18637/jss.v082.i13.

Laine VN, Gossmann TI, Schachtschneider KM, Garroway CJ, Madsen O, Verhoeven KJ, De Jager V, Megens H-J, Warren WC, Minx P (2016) Evolutionary signals of selection on cognition from the great tit genome and methylome. Nature Communications 7, 1–9. https://doi.org/10.1038/ncomms10474.

Lamichhaney S, Berglund J, Almén MS, Maqbool K, Grabherr M, Martinez-Barrio A, Promerová M, Rubin C-J, Wang C, Zamani N (2015) Evolution of Darwin/’s finches and their beaks revealed by genome sequencing. Nature 518, 371–375. https://doi.org/10.1038/nature14181.

Lee KA (2006) Linking immune defenses and life history at the levels of the individual and the species. Integrative and Comparative Biology 46, 1000–1015. https://doi.org/10.1093/icb/icl049.

Lee JW, Beebe K, Nangle LA, Jang J, Longo-Guess CM, Cook SA, Davisson MT, Sundberg JP, Schimmel P, Ackerman SL (2006) Editing-defective tRNA synthetase causes protein misfolding and neurodegeneration. Nature 443, 50–55. https://doi.org/10.1038/nature05096.

Leroy T, Anselmetti Y, Tilak M-K, Bérard S, Csukonyi L, Gabrielli M, Scornavacca C, Milá B, Thébaud C, Nabholz B (2021) A bird’s white-eye view on avian sex chromosome evolution. Peer Community Journal 1, e63. https://doi.org/10.24072/pcjournal.70.

Leroy T, Rousselle M, Tilak M-K, Caizergues AE, Scornavacca C, Recuerda M, Fuchs J, Illera JC, De Swardt DH, Blanco G (2021) Island songbirds as windows into evolution in small populations. Current Biology 31, 1303–1310.e4. https://doi.org/10.1016/j.cub.2020.12.040.

Levin II, Zwiers P, Deem SL, Geest EA, Higashiguchi JM, Iezhova TA, Jiménez-Uzcátegui G, Kim DH, Morton JP, Perlut NG, Renfrew RB, Sari EHR, Valkiunas G, Parker PG (2013) Multiple Lineages of Avian Malaria Parasites (Plasmodium) in the Galapagos Islands and Evidence for Arrival via Migratory Birds. Conservation Biology 27, 1366–1377. https://doi.org/10.1111/cobi.12127.

Levy H, Fiddaman SR, Vianna JA, Noll D, Clucas GV, Sidhu JK, Polito MJ, Bost CA, Phillips RA, Crofts S (2020) Evidence of pathogen-induced immunogenetic selection across the large geographic range of a wild seabird. Molecular Biology and Evolution 37, 1708–1726. https://doi.org/10.1093/molbev/msaa040.

Li H (2013) Aligning sequence reads, clone sequences and assembly contigs with BWA-MEM. ArXiv Preprint ArXiv:13033997. https://doi.org/10.48550/arXiv.1303.3997

Li H, Handsaker B, Wysoker A, Fennell T, Ruan J, Homer N, Marth G, Abecasis G, Durbin R (2009) The Sequence Alignment/Map format and SAMtools. Bioinforma Oxf Engl 25, 2078–2079. https://doi.org/10.1093/bioinformatics/btp352.

Lindström KM, Foufopoulos J, Pärn H, Wikelski M (2004) Immunological investments reflect parasite abundance in island populations of Darwin’s finches. Proceedings of the Royal Society of London Series B: Biological Sciences 271, 1513–1519. https://doi.org/10.1098/rspb.2004.2752.

Lobato E, Doutrelant C, Melo M, Reis S, Covas R (2017) Insularity effects on bird immune parameters: A comparison between island and mainland populations in West Africa. Ecology and Evolution 7, 3645–3656. https://doi.org/10.1002/ece3.2788.

Loire E, Chiari Y, Bernard A, Cahais V, Romiguier J, Nabholz B, Lourenço JM, Galtier N (2013) Population genomics of the endangered giant Galapagos tortoise. Genome Biology 14, R136. https://doi.org/10.1186/gb-2013-14-12-r136.

Loiseau C, Melo M, Lobato E, Beadell JS, Fleischer RC, Reis S, Doutrelant C, Covas R (2017) Insularity effects on the assemblage of the blood parasite community of the birds from the Gulf of Guinea. Journal of Biogeography 44, 2607–2617. https://doi.org/10.1111/jbi.13060

Lomolino MV (2005) Body size evolution in insular vertebrates: generality of the island rule. Journal of Biogeography 32, 1683–1699. https://doi.org/10.1111/j.1365-2699.2005.01314.x.

Losos JB, Ricklefs RE (2009) Adaptation and diversification on islands. Nature 457, 830–836. https://doi.org/10.1038/nature07893.

Lundberg M, Liedvogel M, Larson K, Sigeman H, Grahn M, Wright A, Åkesson S, Bensch S (2017) Genetic differences between willow warbler migratory phenotypes are few and cluster in large haplotype blocks. Evolution Letters 1, 155–168. https://doi.org/10.1002/evl3.15.

Luo R, Liu B, Xie Y, Li Z, Huang W, Yuan J, He G, Chen Y, Pan Q, Liu Y (2012) SOAPdenovo2: an empirically improved memory-efficient short-read de novo assembler. Gigascience 1,. https://doi.org/10.1186/2047-217X-1-18.

MacArthur RH, Wilson EO (1967) The theory of island biogeography. ‘Theory Isl. Biogeogr.’ (Princeton university press)

Maria L, Svensson E, Ricklefs RE (2009) Low diversity and high intra-island variation in prevalence of avian Haemoproteus parasites on Barbados, Lesser Antilles. Parasitology 136, 1121–1131. https://doi.org/10.1017/S0031182009990497.

Martinez J, Vasquez RA, Venegas C, Merino S (2015) Molecular characterisation of haemoparasites in forest birds from Robinson Crusoe Island: is the Austral Thrush a potential threat to endemic birds? Bird Conservation International 25, 139–152. https://doi.org/10.1017/S0959270914000227.

Matson KD (2006) Are there differences in immune function between continental and insular birds? Proceedings Biological Sciences / The Royal Society 273, 2267–2274. https://doi.org/10.1098/rspb.2006.3590.

Matson KD, Beadell JS (2010) Infection, immunity, and island adaptation in birds.

Minias P, Pikus E, Whittingham LA, Dunn PO (2019) Evolution of copy number at the MHC varies across the avian tree of life. Genome Biology and Evolution 11, 17–28. https://doi.org/10.1093/gbe/evy253.

Mueller JC, Kuhl H, Timmermann B, Kempenaers B (2016) Characterization of the genome and transcriptome of the blue tit Cyanistes caeruleus: polymorphisms, sex-biased expression and selection signals. Molecular Ecology Resources 16, 549–561. https://doi.org/10.1111/j.1944-9720.1986.tb01032.x.

Munoz-Mérida A, Viguera E, Claros MG, Trelles O, Pérez-Pulido AJ (2014) Sma3s: a three-step modular annotator for large sequence datasets. DNA Research 21, 341–353. https://doi.org/10.1093/dnares/dsu001.

Nguyen L-T, Schmidt HA, Haeseler A, Minh BQ (2014) IQ-TREE: a fast and effective stochastic algorithm for estimating maximum-likelihood phylogenies. Molecular Biology and Evolution 32, 268–274. https://doi.org/10.1093/molbev/msu300.

Nieberding C, Morand S, Libois R, Michaux J (2006) Parasites and the island syndrome: the colonization of the western Mediterranean islands by Heligmosomoides polygyrus (Dujardin, 1845. Journal of Biogeography 33, 1212–1222. https://doi.org/10.1111/j.1365-2699.2006.01503.x.

O’Connor EA, Hasselquist D, Nilsson J-Å, Westerdahl H, Cornwallis CK (2020) Wetter climates select for higher immune gene diversity in resident, but not migratory, songbirds. Proceedings of the Royal Society B: Biological Sciences 287, 20192675. https://doi.org/10.1098/rspb.2019.2675.

Ohta T (1992) The nearly neutral theory of molecular evolution. Annual Review of Ecology and Systematics 23, 263–286. http://www.jstor.org/stable/2097289.

Ortutay C, Vihinen M (2009) Identification of candidate disease genes by integrating Gene Ontologies and protein-interaction networks: case study of primary immunodeficiencies. Nucleic Acids Research 37, 622–628. https://doi.org/10.1093/nar/gkn982.

Padilla DP, Illera JC, Gonzalez-Quevedo C, Villalba M, Richardson DS (2017) Factors affecting the distribution of haemosporidian parasites within an oceanic island. International Journal for Parasitology 47, 225–235. https://doi.org/10.1016/j.ijpara.2016.11.008.

Paradis E, Schliep K (2019) ape 5.0: an environment for modern phylogenetics and evolutionary analyses in R. Bioinformatics 35, 526–528. https://doi.org/10.1093/bioinformatics/bty633.

Peona V, Blom MPK, Xu L, Burri R, Sullivan S, Bunikis I, Liachko I, Haryoko T, Jønsson KA, Zhou Q (2021) Identifying the causes and consequences of assembly gaps using a multiplatform genome assembly of a bird-of-paradise. Molecular ecology resources 21, 263–286. https://doi.org/10.1111/1755-0998.13252.

Peona V, Weissensteiner MH, Suh A (2018) How complete are “complete” genome assemblies?—An avian perspective. Molecular ecology resources 18, 1188–1195. https://doi.org/10.1111/1755-0998.12933.

Pérez-Rodríguez A, Ramírez Á, Richardson DS, Pérez-Tris J (2013) Evolution of parasite island syndromes without long-term host population isolation: Parasite dynamics in Macaronesian blackcaps Sylvia atricapilla. Global Ecology and Biogeography 22, 1272–1281. https://doi.org/10.1111/geb.12084.

Pinheiro J, Bates D, DebRoy S, Sarkar D, Heisterkamp S, Willigen B, Maintainer R (2017) Package ‘nlme. Linear Nonlinear Mix Eff Models Version 3,. https://svn.r-project.org/R-packages/trunk/nlme/.

R Core Team (2018) R: A language and environment for statistical computing.

Rando JC, Alcover JA, Illera JC (2010) Disentangling Ancient Interactions: A New Extinct Passerine Provides Insights on Character Displacement among Extinct and Extant Island Finches. PLoS One 5:e12956,. https://doi.org/10.1371/journal.pone.0012956.

Ranwez V, Harispe S, Delsuc F, Douzery EJ (2011) MACSE: Multiple Alignment of Coding SEquences accounting for frameshifts and stop codons. PLoS One 6:e22594,. https://doi.org/10.1371/journal.pone.0022594.

Recuerda M, Vizueta J, Cuevas-Caballé C, Blanco G, Rozas J, Milá B (2021) Chromosome-level genome assembly of the common chaffinch (Aves: Fringilla coelebs): a valuable resource for evolutionary biology. Genome Biology and Evolution 13, evab034. https://doi.org/10.1093/gbe/evab034.

Robinson JA, Ortega-Del Vecchyo D, Fan Z, Kim BY, Marsden CD, Lohmueller KE, Wayne RK (2016) Genomic flatlining in the endangered island fox. Current Biology 26, 1183–1189. https://doi.org/10.1016/j.cub.2016.02.062.

Rogers RL, Slatkin M (2017) Excess of genomic defects in a woolly mammoth on Wrangel island. PLoS Genet 13:e1006601,. https://doi.org/10.1371/journal.pgen.1006601.

Rohland N, Reich D (2012) Cost-effective, high-throughput DNA sequencing libraries for multiplexed target capture. Genome Research 22, 939–946. https://doi.org/10.1101/gr.128124.111.

Rousselle M, Simion P, Tilak M-K, Figuet E, Nabholz B, Galtier N (2020) Is adaptation limited by mutation? A timescale-dependent effect of genetic diversity on the adaptive substitution rate in animals. PLoS Genetics 16, e1008668. https://doi.org/10.1371/journal.pgen.1008668.

Santonastaso T, Lighten J, Oosterhout C, Jones KL, Foufopoulos J, Anthony NM (2017) The effects of historical fragmentation on major histocompatibility complex class II β and microsatellite variation in the Aegean island reptile, Podarcis erhardii. Ecology and Evolution 7, 4568–4581. https://doi.org/10.1002/ece3.3022.

She R, Chu JS-C, Uyar B, Wang J, Wang K, Chen N (2011) genBlastG: using BLAST searches to build homologous gene models. Bioinformatics 27, 2141–2143. https://doi.org/10.1093/bioinformatics/btr342.

Shultz AJ, Sackton TB (2019) Immune genes are hotspots of shared positive selection across birds and mammals. Elife 8, e41815. https://doi.org/10.7554/eLife.41815.

Siewert KM, Voight BF (2020) BetaScan2: Standardized Statistics to Detect Balancing Selection Utilizing Substitution Data. Genome Biology and Evolution 12, 3873–3877. https://doi.org/10.1093/gbe/evaa013.

Silva-Iturriza A, Ketmaier V, Tiedemann R (2012) Prevalence of avian haemosporidian parasites and their host fidelity in the central Philippine islands. Parasitology International 61, 650–657. https://doi.org/10.1016/j.parint.2012.07.003.

Simion P, Belkhir K, François C, Veyssier J, Rink JC, Manuel M, Philippe H, Telford MJ (2018) A software tool ‘CroCo’detects pervasive cross-species contamination in next generation sequencing data. BMC Biology 16, 1–9. https://doi.org/10.1186/s12915-018-0486-7.

Singhal S, Leffler EM, Sannareddy K, Turner I, Venn O, Hooper DM, Strand AI, Li Q, Raney B, Balakrishnan CN (2015) Stable recombination hotspots in birds. Science 350, 928–932. https://doi.org/10.1126/science.aad0843.

Slade RW, McCallum HI (1992) Overdominant vs. frequency-dependent selection at MHC loci. Genetics 132,. https://doi.org/10.1093%2Fgenetics%2F132.3.861.

Slowikowski K, Schep A, Hughes S, Lukauskas S, Irisson J-O, Kamvar ZN, Ryan T, Christophe D, Hiroaki Y, Gramme P (2018) Package ggrepel. Autom Position Non-Overlapping Text Labels ggplot2,.

Smeds L, Qvarnstrom A, Ellegren H (2016) Direct estimate of the rate of germline mutation in a bird. Genome Research gr-204669. https://doi.org/10.1101/gr.204669.116.

Spiess A-N, Spiess MA-N (2018) Package ‘qpcR. ‘Model Anal Real-Time PCRdata Httpscran R-Proj’. (OrgwebpackagesqpcRqpcR Pdf)

Spurgin LG, Van Oosterhout C, Illera JC, Bridgett S, Gharbi K, Emerson BC, Richardson DS (2011) Gene conversion rapidly generates major histocompatibility complex diversity in recently founded bird populations. Molecular Ecology 20, 5213–5225. https://doi.org/10.1111/j.1365-294X.2011.05367.x

Tange O (2018) GNU parallel 2018. https://doi.org/10.5281/zenodo.1146014.

Van Riper III C, Van Riper SG, Goff ML, Laird M (1986) The epizootiology and ecological significance of malaria in Hawaiian land birds. Ecological Monographs 56, 327–344. https://doi.org/10.2307/1942550.

Velová H, Gutowska-Ding MW, Burt DW, Vinkler M (2018) Toll-like receptor evolution in birds: gene duplication, pseudogenization, and diversifying selection. Mol Biol Evol 35, 2170–2184. https://doi.org/10.1093/molbev/msy119

Warren WC, Clayton DF, Ellegren H, Arnold AP, Hillier LW, Künstner A, Searle S, White S, Vilella AJ, Fairley S (2010) The genome of a songbird. Nature 464, 757–762. https://doi.org/10.1038/nature08819.

Warren BH, Simberloff D, Ricklefs RE, Aguilée R, Condamine FL, Gravel D, Morlon H, Mouquet N, Rosindell J, Casquet J (2015) Islands as model systems in ecology and evolution: Prospects fifty years after MacArthur-Wilson. Ecology Letters 18, 200–217. https://doi.org/10.1111/ele.12398.

Welch JJ, Eyre-Walker A, Waxman D (2008) Divergence and Polymorphism Under the Nearly Neutral Theory of Molecular Evolution. Journal of Molecular Evolution 67, 418–426. https://doi.org/10.1007/s00239-008-9146-9.

Wickham H (2016) ggplot2: Elegant Graphics for Data Analysis. https://doi.org/10.1007/978-3-319-24277-4_9.

Wikelski M, Foufopoulos J, Vargas H, Snell H (2004) Galápagos birds and diseases: invasive pathogens as threats for island species. Ecology and Society 9,. http://www.jstor.org/stable/26267654.

Wolf JBW, Künstner A, Nam K, Jakobsson M, Ellegren H (2009) Nonlinear Dynamics of Nonsynonymous (dN) and Synonymous (dS) Substitution Rates Affects Inference of Selection. Genome Biol Evol 1, 308–319. https://doi.org/10.1093/gbe/evp030.

Zhang G, Parker P, Li B, Li H, Wang J (2012) The genome of Darwin’s Finch (Geospiza fortis). 297 MB. https://doi.org/10.5524/100040.

