## Supplementary material for "Evolution of immune genes in island birds: reduction in population sizes can explain island syndrome": https://doi.org/10.6084/m9.figshare.16954921.v7

Peer Community In Evolutionary Biology

RESEARCH ARTICLE
Open Access
Open Data
Open Code
Open Peer-Review

### Supplementary materials for "Evolution of immune genes in island birds: reduction in population sizes can explain island syndrome."

Mathilde BARTHE[1*], Claire DOUTRELANT[2], Rita COVAS[3,5], Martim MELO[3-6], Juan Carlos ILLERA[7], Marie-Ka TILAK[1], Constance COLOMBIER[1], Thibault LEROY[1,8], Claire LOISEAU[2,3,4a], Benoit NABHOLZ[1,9a]

**Cite as**
Barthe M, Nabholz B (2022) Supplementary materials for "Evolution of immune genes in island birds: reduction in population sizes can explain island syndrome." Figshare. https://doi.org/10.6084/m9.figshare.16954921

**Correspondence**


**Recommender**
Emma Berdan

**Reviewers**
Steven Fiddaman and three anonymous reviewers

#### Supplementary Methods :

##### Test for contamination

Analyses were performed to identify contaminations between individuals of different species. First, we generate assembly for all individuals using Megahits v.1.2.7 (Li et al. 2015). Next, we use the program CroCo (vers. 1.1; Simion et al. 2018) to identify candidates for cross-species contamination. CroCo identifies a contig from the assembly of the species X as contaminated by the species Y if the reads from the species Y map with a higher relative proportion on the contig than the reads for the species X.

Contamination tests were performed on 3765 combinations of individuals from different species. On average, 3.12% of the contigs were considered as contaminants (Figure S1). The top 20% of the combinations (i.e., a value greater than 3.09% of contaminating contigs ; Figure S1) always involve a pair of species belonging to the same genus. In this case, contamination could be difficult to identify due to the low genetic divergence between species. The remaining 80% showed low percentages of contamination (<5% of scaffolds) that may be associated with false positives (i.e., corresponding to highly conserved sequences between species). Overall, we did not detect a clear case of cross-species contamination in our dataset.

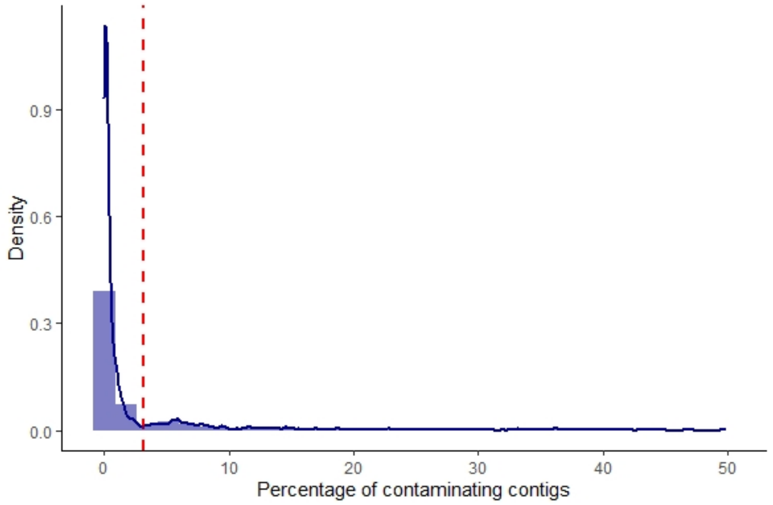


**Figure S1:** Distribution of the percentage of contaminating contigs. The red line represents the 80% quantile.

**Homology identification**

After defining the genes composition of our differents group, see Selection and identification of immune and control genes, we obtained their reference sequences from the Ensembl database for core groupe and control group and from the annotated genome of *F. albicolis for* Database groupe and Sma3s groupe. We also used the TLRs alignments from Velová et al. (2018).

Then, in order to identify corresponding sequences in our 11 reference genomes, we search for homologous sequences. To do so, we first translated sequences with transeq (vers. 6.6.0.0; Rice et al. 2000), then we used the program Silix (vers. 1.2.9; Miele et al. 2011) to cluster amino-acid sequences with at least 60% of similarity and a minimum overlap of 30%. For partial sequences, a minimum length of 10 amino acids or a minimum of 5% of the complete protein sequence was required.

For the immunity genes, some sequences could be assigned to several clusters due to paralogy (*e.g.*, TLR2A and TLR2B). In the core group, we use the probabilistic approach of the profile hidden Markov models implemented in HMMER (v. 3.3.2; Mistry et al. 2013) to assign the gene to a cluster. We first created a gene-specific HMM profile from the alignments of Velová et al. (2018). Then, the candidate sequences were assigned to each homologous group based on the best score of a hmmscan search. For larger sets of genes (i.e Database-group and Sma3s-group), we remove duplicated sequences in the dataset to analyse only one.

**Detection of genes under balancing selection**

The Database-group and Sma3s-group most likely include immune genes subject to different selection pressures (*i.e.*, balancing and purifying selection). To identify genes evolving under balancing selection, unfolded SNP frequency was estimated using a home-made script for *Taeniopygia guttata* using *Poephila acuticauda* as outgroup. The allelic state of *Poephila* was taken as the ancestral state of *Taeniopygia*. Then, we run the program BetaScan (vers. 2; Siewert and Voight 2020) with a sliding window of 2000pb and option '-fold -m 0.15' allowed to exclude allele with a frequency under 15% to avoid false positives to identify SNPs. We selected the top SNPs with a score above a 99.9% threshold based on the Beta* score as evolving under balancing selection (Siewert and Voight 2020). Finally, genes overlapping with the top SNPs were considered as genes evolving under balancing selection or overdominance. Respectively only 2 and 3 genes from Database-group and Sma3s-group sets were identified and removed from the analysis.

**Simulation of control genes under balancing selection**

Simulations were performed using SLiM software (vers. 3.3.2; Haller and Messer 2017). Sequences of 300kb with a mutation of 4.6e-9 substitutions/site/generation were simulated (Smeds et al. 2016). Recombination was set to be equal to mutation rate. Three types of mutations were possible: i) neutral synonymous mutations, ii) non-synonymous mutations with a Distribution of Fitness Effect (DFE) following a gamma law of mean = -0.025 and shape = 0.3, which corresponds to the DFE estimated in Passerines by Rousselle et al. (2020), iii) non-synonymous mutations under balancing selection with an effect on fitness initially set at 0.01 but re-estimated by the program at each generation according to the mutation frequency in the population, thus including a frequency-dependent effect.

We simulate a coding sequence organization where positions one and two of the codons were considered as non-degenerated sites with the two non-synonymous types of mutations previously described are possible in proportion of 1 mutation under balancing selection for 6 randomly sampled in the DFE. The third position was considered as a 4-fold degenerate site, where only synonymous mutations could appear.

Two population sizes were simulated. One of 270,000 individuals representing large mainland population sizes, and another of 110,000 individuals representing smaller island populations. These values were estimated from the synonymous nucleotide diversity (Ps) estimated for insular and mainland species in our dataset (see part *Polymorphism analyses* from Materials) which allow the estimation of the effective population size as Ne = Ps / 4*µ (with µ=4.6e-9 corresponding to mutation rate estimated for the collared flycatcher in Smeds et al. 2016). Simulations were replicated 15 times, by sampling the genome of 15 individuals after 100,000 generations, of which 10,000 generations corresponded to a burning phase. Nucleotide diversity of non-degenerated and 4-fold degenerate sites were estimated by arithmetic averaging the divergences of all sequence pairs (Tajima 1989).

Next, we simulated the effect of population size variation on Pn/Ps under two types of balancing selection, namely frequency-dependence and overdominance (Figure S4, S5). In both cases, we simulated 10 population sizes for both island and mainland populations (starting respectively from 1100 and 2700 individuals and doubled between each population size step), 10 replicas were made for each population size. We use the same parameters as before, except for the non-synonymous mutation, where for i) frequency dependence, mutations were initially set a fitness effect of 0.01 and then re-estimated at each generation according to the mutation frequency in the population, and ii) for overdominance, mutations sets a fitness effect of 0.01 and a dominance coefficient of 1.5.

Finally, we tested the effect of selection coefficient variation. To do so, we fixed population size at 110,000 individuals and made the fitness vary from 0 to 0.045 for frequency dependence mutation (Figure S6A), and the dominance coefficient (h) from 1.0 to 1.8 for overdominance mutations (Figure S6B).

#### Supplementary figures and tables:

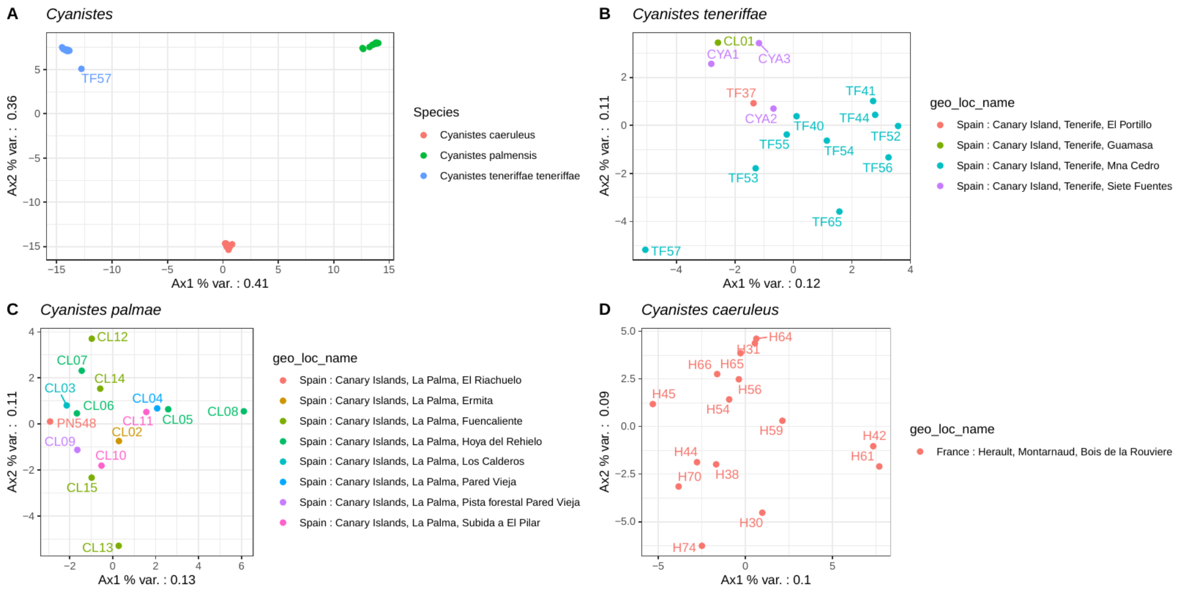


**Figure S2:** Principal Component Analysis PCA using alleles frequencies of the control genes. A) All *Cyanistes* species; B) *Cyanistes teneriffae*; C) *Cyanistes palmensis*; D) *Cyanistes caeruleus*. Colors indicate species or locality.

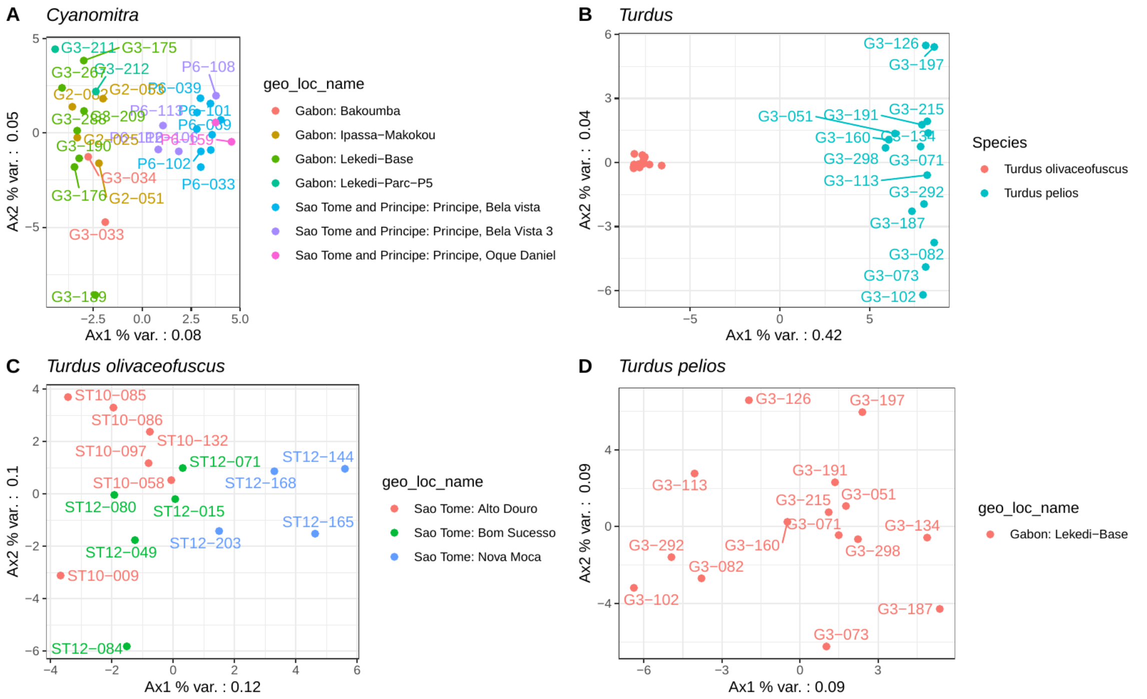


**Figure S3:** Principal Component Analysis PCA using alleles frequencies of the control genes. A) *Cyanomitra olivacea;* B) All *Turdus* species; C) *Turdus olivaceofuscus*; D) *Turdus pelios*. Colors indicate species or locality.

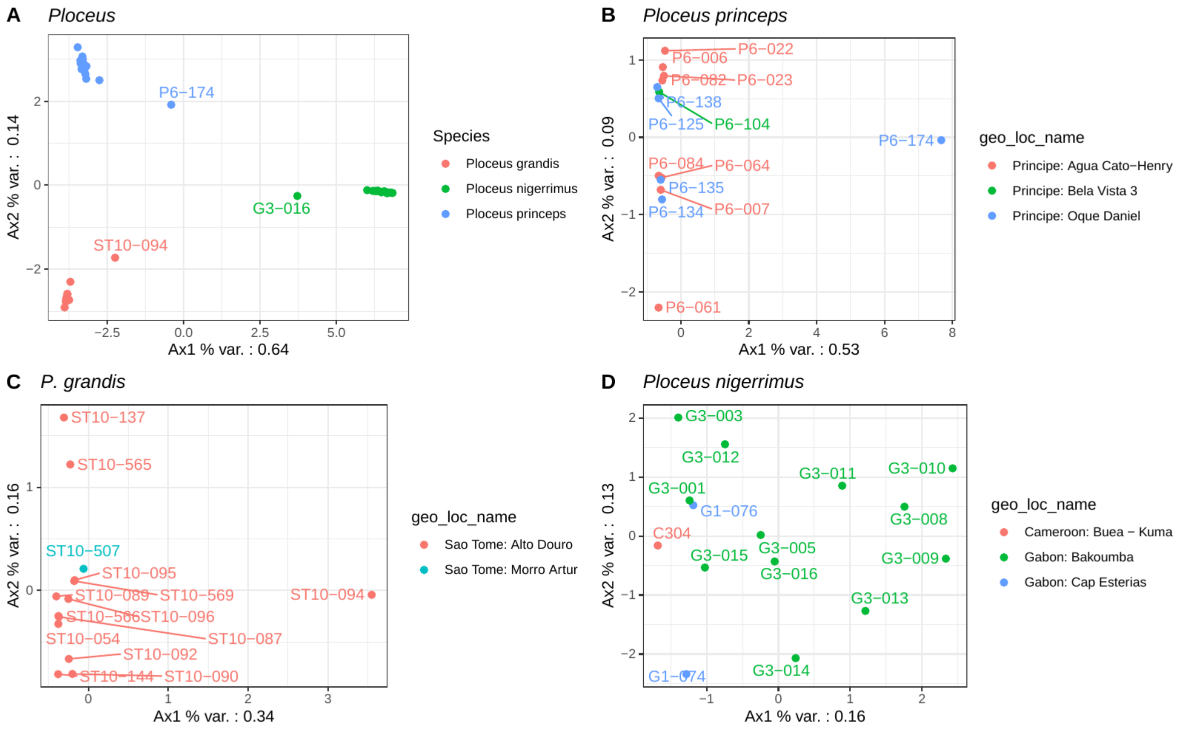


**Figure S4:** Principal Component Analysis PCA using alleles frequencies of the control genes. A) All *Ploceus* species; B) *Ploceus princeps*; C) *Ploceus grandis*; D) *Ploceus nigerrimus*. Colors indicate species or locality.

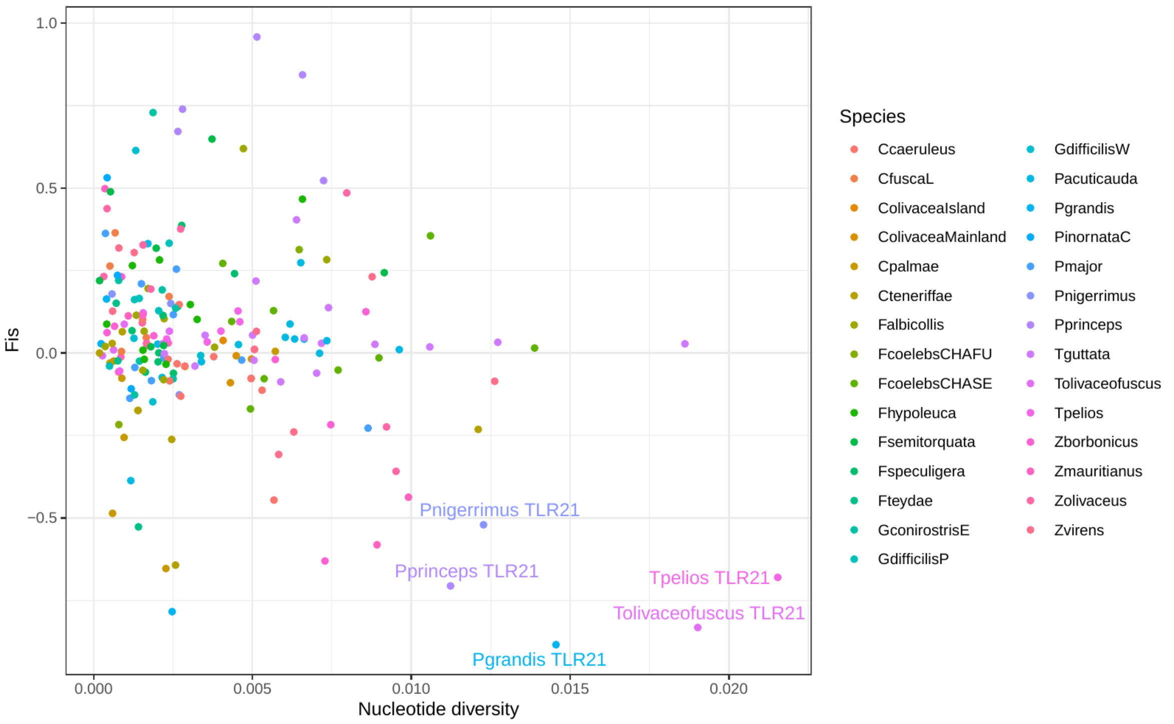


**Figure S5:** Deviation of heterozygosity from Hardy-Weinberg proportion (Fis) according to nucleotide diversity (Pi). A negative Fis indicates more heterozygous individuals than expected.

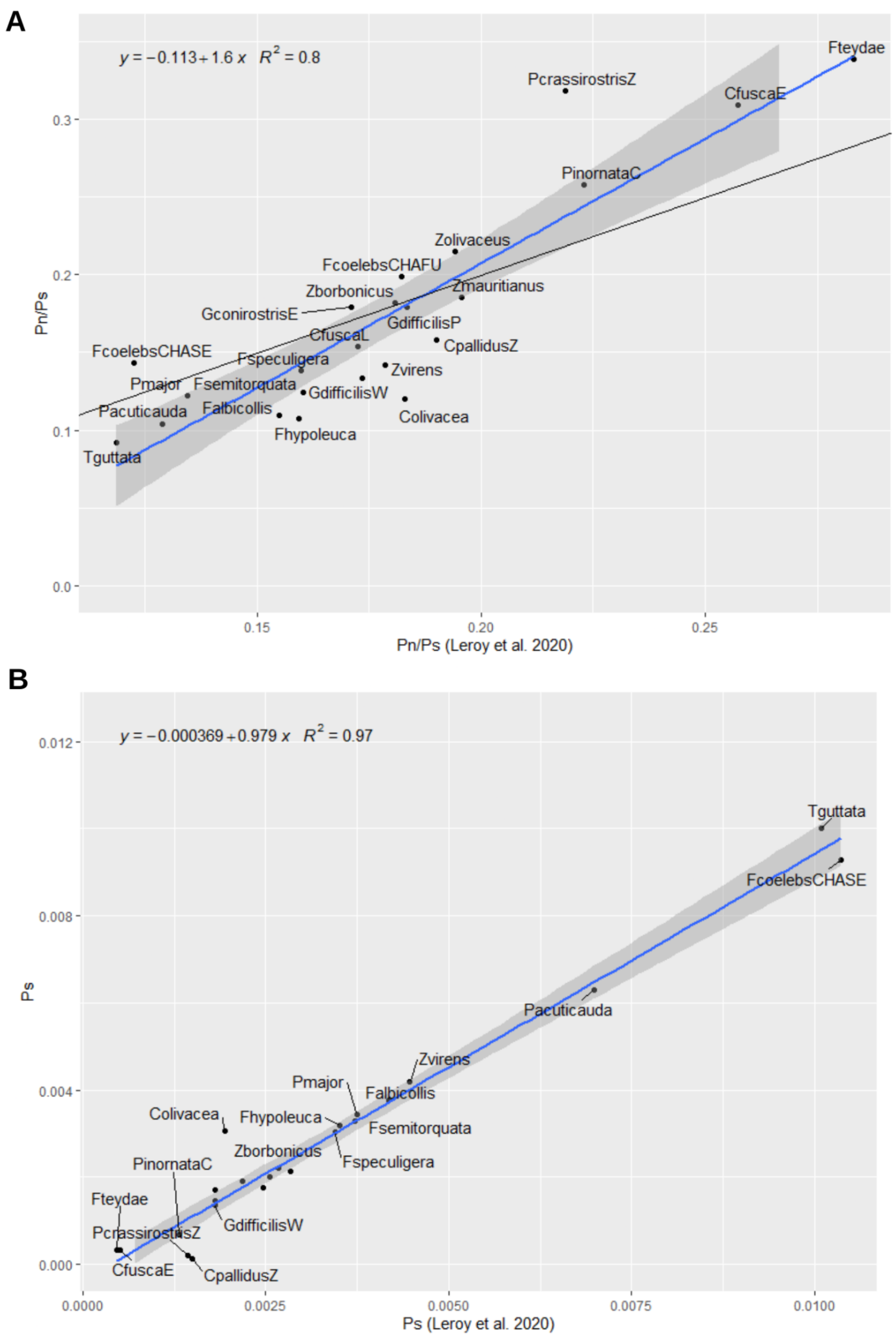


**Figure S6:** Correlation between Pn/Ps (a) and Ps (b) calculated on the control genes in this study's dataset and those calculated by Leroy et al. (2021). Ps: synonymous polymorphism Pn: non-synonymous polymorphism.

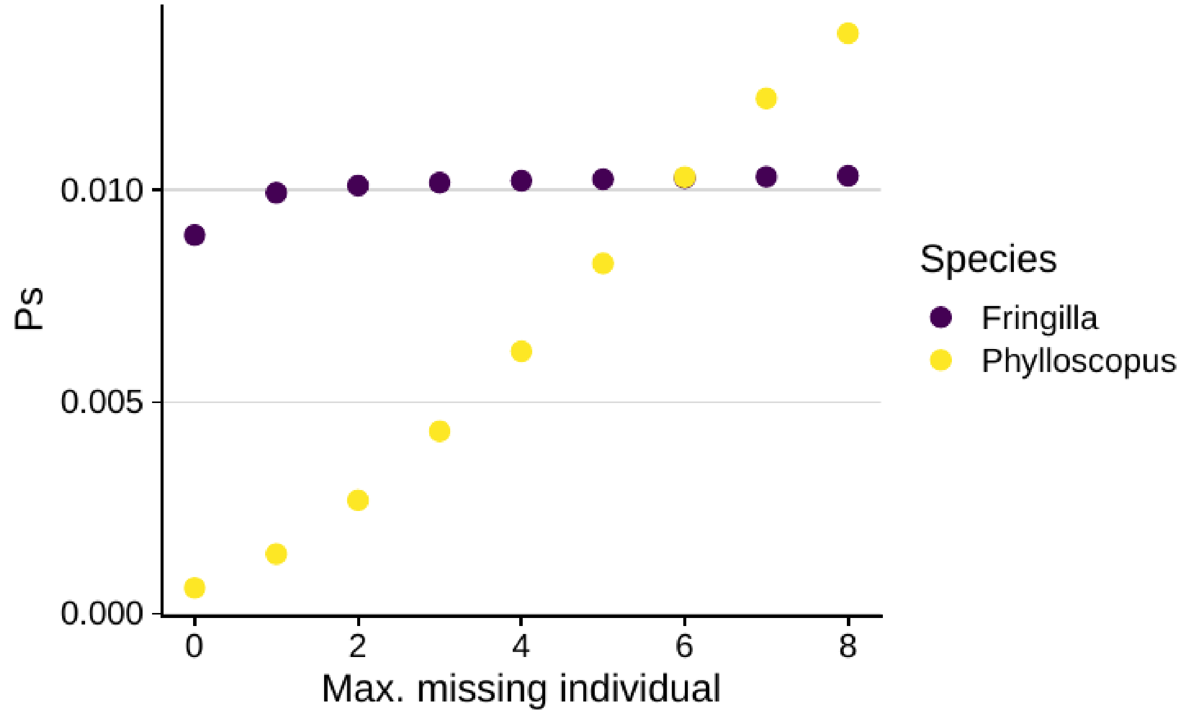


**Figure S7**: Relationship between the maximum number of missing individuals allowed and synonymous nucleotide diversity (Ps)in *Phylloscopus trochilus* and *Fringilla coelebs*.

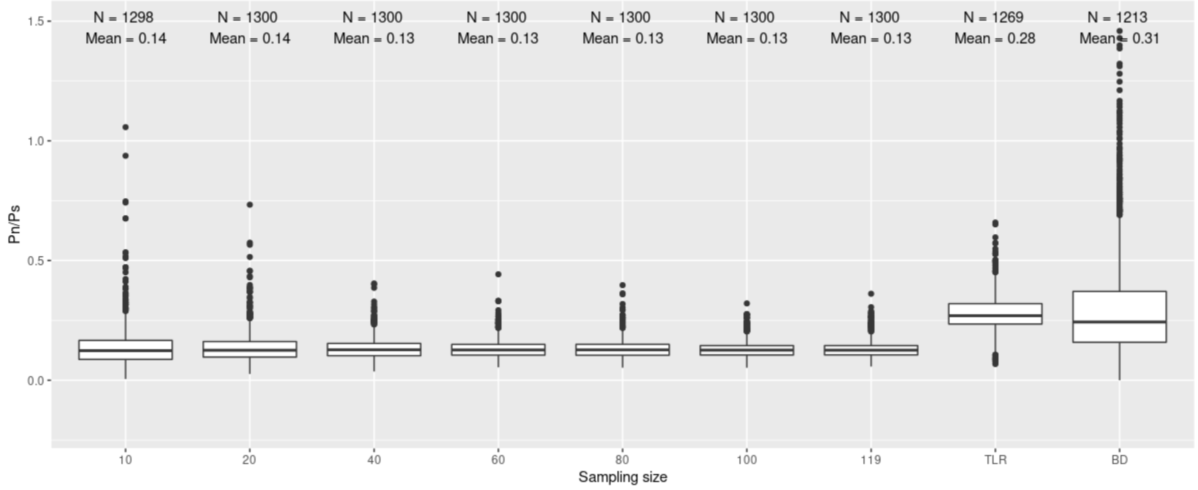


**Figure S8**: Effect of sub-sampling size on PN/PS. Bootstrap of 100.For BD samplig of 9 genes, for TLR sampling of 10 genes.

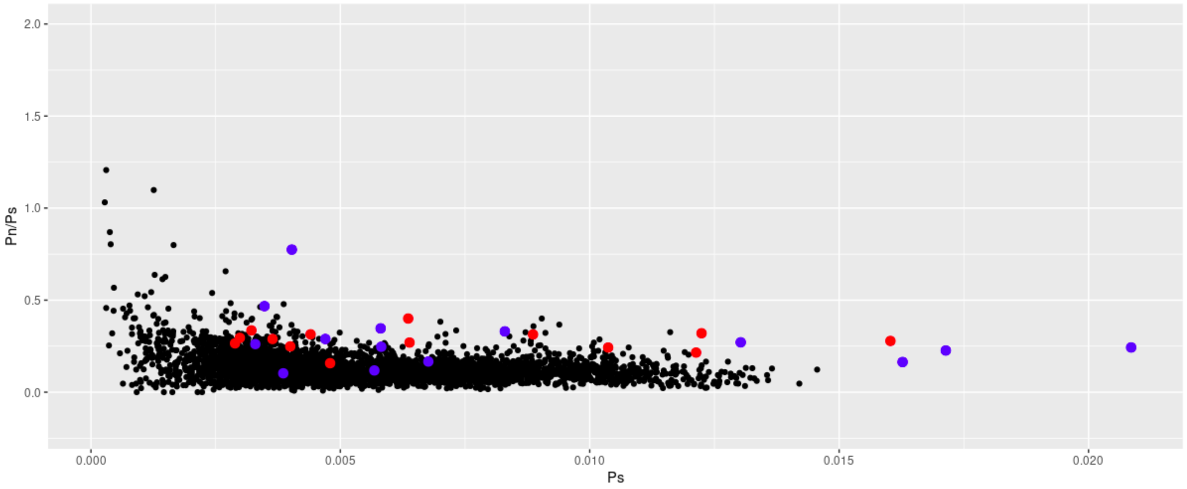


**Figure S9**: Effect of sub-sampling of control genes, here each point corresponds to the values of a mainland species calculated on 10 randomly selected control genes. In blue the BD, in red the TLR gene.

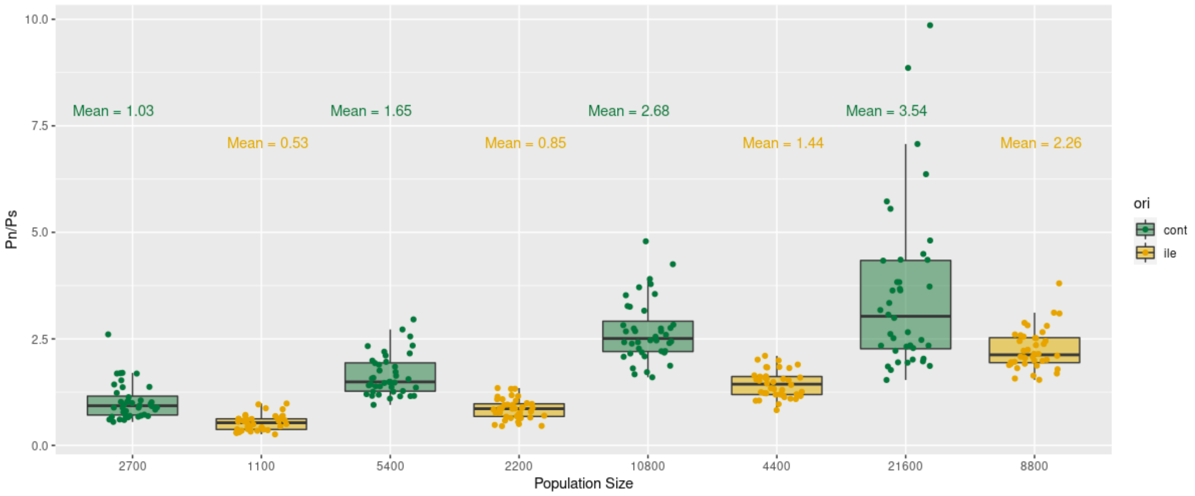


**Figure S10**: Boxplot of Pn/Ps according to population size for simulated sequences under overdominance with SLiM. Dominance coefficient is fixed at 1.5. Each modality is replicated 10 times.

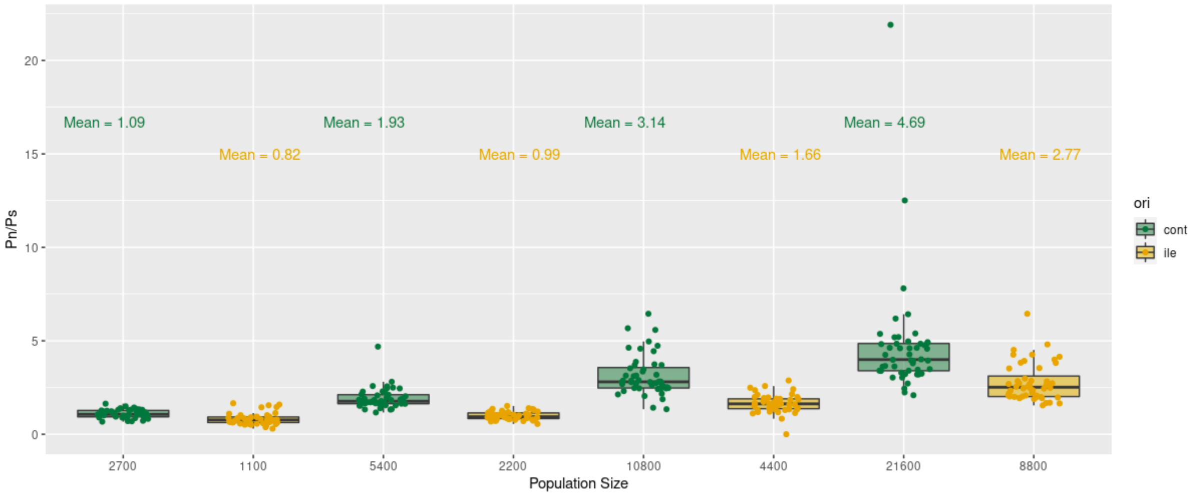


**Figure S11:** Boxplot of Pn/Ps according to population size for simulated sequences under frequency dependence with SLiM. Each modality is replicated 10 times.

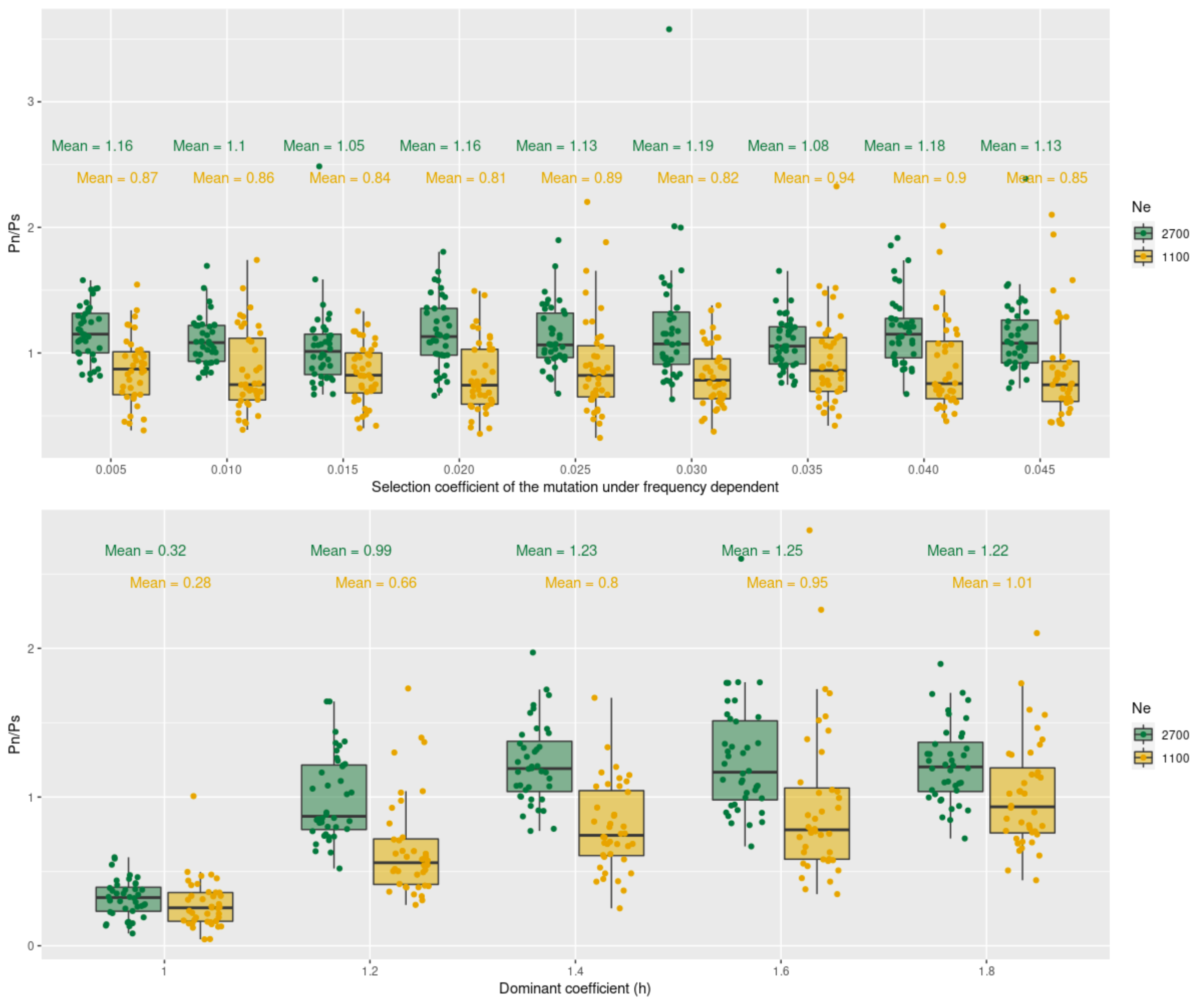


**Figure S12:** Boxplot of Pn/Ps according to a) initial selection coefficient of the mutation under frequency dependence b) dominance coefficient (h) for simulated sequences under overdominance with SLiM. Each modality is replicated 10 times.

**Table S1:** Model selection of all gene categories using a reduced number of families (we grouped Turdidae within Muscicapidae, Nectariniidae, and Estrildidae within Ploceidae and Fringillidae within Thraupidae).
https://figshare.com/s/ab7004cc2f4415b4058f

**Table S2**: Summary of the linear model and PGLS model using phylogeny from Figure 1 and the *Δ*Pn/Ps ( difference between the Pn/Ps of immune genes and control genes).
https://figshare.com/s/ab7004cc2f4415b4058f

**Table S3**: Table with information regarding the samples newly-sequenced in this study.
https://figshare.com/s/ab7004cc2f4415b4058f

**Table S4**: Table with information regarding the sample obtained from Leroy et al. 2021.
https://figshare.com/s/ab7004cc2f4415b4058f

**Table S5 to S14:** Model selection by AICc criterion and ANOVA test. Summary of the best models.
https://figshare.com/s/ab7004cc2f4415b4058f

**Table S15 to S24:**
https://figshare.com/s/ab7004cc2f4415b4058f

#### References

Barthe M, Doutrelant C, Covas R, Melo M, Illera JC, Tilak M-K, Colombier C, Leroy T, Loiseau C, Nabholz B (2022) Evolution of immune genes in island birds: reduction in population sizes can explain island syndrome. bioRxiv, 2021.11.21.469450, ver. 4 peer-reviewed and recommended by Peer Community in Evolutionary Biology. https://doi.org/10.1101/2021.11.21.469450

Haller BC, Messer PW (2017) SLiM 2: Flexible, interactive forward genetic simulations. *Molecular Biology and Evolution* **34**, 230–240. doi:https://doi.org/10.1093/molbev/msw211

Li D, Liu C-M, Luo R, Sadakane K, Lam T-W (2015) MEGAHIT: an ultra-fast single-node solution for large and complex metagenomics assembly via succinct de Bruijn graph. *Bioinformatics* **31**, 1674–1676. doi:https://doi.org/10.1093/bioinformatics/btv033

Miele V, Penel S, Duret L (2011) Ultra-fast sequence clustering from similarity networks with SiLiX. *BMC Bioinformatics* **12**, 1–9. doi:https://doi.org/10.1186/1471-2105-12-116

Mistry J, Finn RD, Eddy SR, Bateman A, Punta M (2013) Challenges in homology search: HMMER3 and convergent evolution of coiled-coil regions. *Nucleic Acids Research* **41**, e121–e121. doi:https://doi.org/10.1093/nar/gkt263

Rice P, Longden I, Bleasby A (2000) EMBOSS: the European molecular biology open software suite. *Trends in Genetics* **16**, 276–277. https://doi.org/10.1016/S0168-9525(00)02024-2

Rousselle M, Simion P, Tilak M-K, Figuet E, Nabholz B, Galtier N (2020) Is adaptation limited by mutation? A timescale-dependent effect of genetic diversity on the adaptive substitution rate in animals. *PLoS Genetics* **16**, e1008668. doi:https://doi.org/10.1371/journal.pgen.1008668

Siewert KM, Voight BF (2020) BetaScan2: Standardized Statistics to Detect Balancing Selection Utilizing Substitution Data. *Genome Biology and Evolution* **12**, 3873–3877. doi:https://doi.org/10.1093/gbe/evaa013

Simion P, Belkhir K, François C, Veyssier J, Rink JC, Manuel M, Philippe H, Telford MJ (2018) A software tool ‘CroCo’detects pervasive cross-species contamination in next generation sequencing data. *BMC Biology* **16**, 1–9. doi:https://doi.org/10.1186/s12915-018-0486-7

Smeds L, Qvarnstrom A, Ellegren H (2016) Direct estimate of the rate of germline mutation in a bird. *Genome Research* gr-204669. doi:https://doi.org/10.1101/gr.204669.116

Tajima F (1989) Statistical method for testing the neutral mutation hypothesis by DNA polymorphism. *Genetics* **123**, 585–595. doi:https://doi.org/10.1093/genetics/123.3.585.

Velová H, Gutowska-Ding MW, Burt DW, Vinkler M, Yeager M (2018) Toll-like receptor evolution in birds: gene duplication, pseudogenisation and diversifying selection. *Molecular Biology and Evolution*. doi:https://doi.org/10.1093/molbev/msy119
